# Spatial analysis of canine glioma reveals heterogeneous intratumoral niches characterized by distinct patterns of cell plasticity and immunosuppression

**DOI:** 10.64898/2026.09.24.754227

**Authors:** Sophie R. Nelissen, Andreas Stephanou, Eric N. Glass, James Hammond, Gena Silver, Praveen Sethupathy, Iwijn de Vlaminck, Andrew D. Miller, Elena A. Demeter

## Abstract

Glioma is a fatal tumor of the central nervous system (CNS) with limited therapeutic options. While immune checkpoint inhibitors (ICI) are gaining traction as a novel alternative treatment for human gliomas (HG), clinical trials have yielded mixed results. A proposed major barrier to ICI therapy in cancer is epithelial-mesenchymal plasticity (EMP)-driven intratumoral immunosuppression. However, studying this mechanism is difficult in *ex vivo* and murine models, which fail to adequately replicate cancer cell-immune cell interactions. Formalin-fixed, paraffin-embedded (FFPE) patient-derived canine glioma (CG) samples represent an attractive alternative approach. Although mechanistic perturbation is unachievable in archival FFPE tissues, CG is notoriously “cold” immunologically and is actively being investigated as a comparative model of HG. Moreover, the immune microenvironment in CG advantageously develops under exposure to environmental conditions directly shared with humans. Yet, it remains unknown whether CGs harbor heterogeneous niches combining varied plastic phenotypes and immunosuppressive populations. Defining the immune landscape of CG in relation to EMP is therefore critical to determining its potential as a comparative model for immunotherapy response in HG. By combining standard histology, immunohistochemistry, and tissue microarray-based spatial transcriptomics, we analyzed 30 low-grade (LGr) and high-grade (HGr) FFPE CG cases (n=10 cases/subtype): astrocytomas (4xLGr, 6xHGr), oligodendrogliomas (1xLGr, 9xHGr), and undefined gliomas (3LxGr, 7xHGr). All CG subtypes harbored heterogeneous immunosuppressive niches associated with varied EMP phenotypes. Astrocytomas exhibited a more-mesenchymal identity correlated with recruitment of tumor-associated macrophages (TAMs). Oligodendrogliomas displayed a strong stem-like signature and uniquely upregulated a *FOXP3/LAG3/SNAI1* triad, positioned directly at the interface of immunosuppression and EMP. Across all CG subtypes, a more-mesenchymal phenotype co-occurred with TAMs, and intratumoral T-cell dysfunction and exhaustion. We conclude that CG replicates the EMP-immunosuppression heterogeneity seen in humans. Importantly, our results emphasize the existence of subtype-specific mechanisms of EMP and immune evasion, underscoring the major relevance of subtyping for translational applications.

## Introduction

Gliomas constitute a heterogeneous class of primary tumors of the central nervous system (CNS) that remain almost invariably fatal [1]. Lately, immunotherapy has drawn interest as a promising alternative to vastly-palliative standard treatment options, such as radical surgical resection and (chemo)radiotherapy [2–5]. However, clinical trials have yielded modest success so far, which is thought to partially pertain to the immunologically “cold” nature of gliomas. Moreover, different glioma types exhibit differential sensitivity to immunotherapy [2,3,5–8].

In humans, the WHO classification of gliomas incorporates both histopathology and molecular diagnostics for diagnosis. Adult-onset diffuse gliomas are accordingly divided into isocitrate dehydrogenase 1 (*IDH1*) and 2 (*IDH2*)-mutant astrocytoma and oligodendroglioma, and *IDH*-wildtype glioblastoma (GBM) [1]. In low-grade *IDH-*mutant gliomas, *IDH1* mutation is intrinsically responsible for sparse CD8^+^ cytotoxic T-cell infiltration by repressing STAT1 production [9]. Besides low T-cell permeability, *IDH*-wildtype gliomas have been demonstrated to host a heterogeneous, strongly immunosuppressive microenvironment [7]. Attempts at using immune checkpoint inhibitors (ICI) to treat low-grade glioma and GBM have consequently yielded mixed curative success so far [2,3]. Yet, there is evidence that using PD-1-targeted checkpoint blockade as a neoadjuvant therapy, alone or combined with other ICI, may effectively prolong the OS of GBM patients [4,5].

Aside from overt anatomical hurdles that might directly curb on-target effects, such as the blood-brain barrier, recent advances have shown that epithelial–mesenchymal plasticity (EMP) is a major impediment to effective immunotherapy in cancer [4,10–12]. EMP is a cell phenotype-altering phenomenon that is tightly regulated by specific transcription factors (*ZEB1/2, SNAI1/2, TWIST1/2*) and transforms epithelial neoplastic cells into “more-mesenchymal” cells residing along a spectrum of intermediate phenotypes [13–15]. More-mesenchymal microenvironments foster hypoxic conditions and accumulate major immunosuppressive factors (TGF-β1), thus have been correlated with sparse immune cell infiltration, effector T-cell exhaustion (PD-1, CTLA-4), and increased recruitment of regulatory T-cells (Tregs) and pro-oncogenic M2-like macrophages (TAMs). These microenvironments arise heterogeneously therefore create an intratumoral patchwork of immunosuppressive and immunopermissive regions with differential sensitivity to treatment. EMP consequently curbs the efficacy of immunotherapy significantly [13,14,16].

This deleterious interplay has been identified in a breadth of carcinomas, including breast cancer, colorectal cancer, and squamous cell carcinoma [13,15,16]. Though glial cells are not epithelial *per se*, recent work has highlighted similar plastic trends more specifically in GBM, which could comparably account for heterogeneous patient response to ICI treatment [11,12,17–20]. Building on the originally described classical, mesenchymal, and proneural GBM phenotypes, Neftel *et al*. have recently defined four plastic GBM cell states that may dynamically co-occur within one tumor agnostic of GBM subtype: neural-progenitor (NPC)-like, oligodendrocyte progenitor (OPC)-like, astrocyte (AC)-like, and mesenchymal (MES)-like [18,21,22]. Some pivotal factors for acquiring a GBM MES-like state, such as upregulation of *VIM*, *SNAI1/2*, and *ZEB1/2*, are identical to those responsible for inducing a more-mesenchymal phenotype through EMP [18]. TAMs have additionally been demonstrated to drive GBM cells toward a MES-like state by secreting oncostatin M (OSM), thereby strengthening the parallels with EMP and immunosuppression [23]. While the cell plasticity-immunosuppression axis has been best characterized in GBM, bulk RNA analysis of low-grade HG has highlighted similar trends [24,25]. Unfortunately, understanding how extensively glioma cell plasticity is involved in immunotherapy failure is experimentally complex, since adequately investigating the dynamics of cell plasticity and immunosuppression requires an immunocompetent host. Though transgenic murine models and engraftable glioma cells implanted in syngeneic immunocompetent mice may provide some insight, major anatomical, physiological, and environmental differences exist between humans and laboratory mice (*Mus musculus*). Spontaneous murine glioma is exquisitely rare, further precluding the use of naturally occurring tumors [26]. Additionally, sequencing and transcriptomic analysis of GBM patient-derived samples suggest that those dynamics may be more complex than originally anticipated in glioma, as acquired resistance to immunotherapy in GBM counterintuitively correlates with phenotypic shift away from the MES-like state [10,27].

As biomedical research is rapidly multiplying efforts to reduce reliance on animal models, an attractive alternative is to leverage archived formalin-fixed, paraffin-embedded (FFPE) canine glioma (CG) samples [28,29]. Though CG and human glioma (HG) have distinct biological characteristics, similarities exist between both [30]. Owing notably to multidisciplinary efforts led by the Comparative Brain Tumor Consortium, specific genetic mutations have been identified in CG that resemble those found in HG (*PDGFRA, PIK3CA, TP53*) [31–33]. By contrast, major differences lie within the grading systems used for CG (low-grade, high-grade), and for HG (grade 1-4 attributed within one tumor type), and within the lack of reliance on molecular identity for diagnosis and treatment in CG [31,34]. Importantly, GBM is not recognized as a tumor type in CG given its intrinsic biological and mutational profiles in humans.

Similar to humans, treatment options for CG are broadly limited to surgery, (chemo)radiotherapy, and adjuvant therapy [35]. With the recent release of a canine-specific immune checkpoint inhibitor (ICI) (Gilvetmab®, anti-PD-1), and novel clinical trials (CAR T-cells, anti-CD200 ICI), immunotherapy is also arising as a realistic clinical approach for CG patients [36,37]. Translationally relevant associations between EMP and immunosuppression have been established in canine and feline mammary cancer [36,38,39]. However, such parallels have not yet been established in CG. Compared to a growing body of literature on TAMs, both T-cell exhaustion and regulatory T-cells remain sparsely explored in CG [40–42].

A specific benefit of utilizing patient-derived FFPE CG samples for translational ICI applications is their occurrence in immunocompetent canine patients, meaning that assembly of the immunological tumor microenvironment (TME) was influenced by environmental factors shared with humans. Similarly to humans, evidence of an immunologically “cold” landscape and of EMP pathways upregulation have already been found separately in CG. High-grade CG have been demonstrated to harbor higher numbers of FoxP3^+^ regulatory T-cells and M2-like macrophages than low-grade CG [40,43]. Regarding immunotherapy, a recent study additionally found that French bulldogs with high-grade glioma treated with anti-CD200 ICI had a worse prognosis than other breeds [35]. Remarkably, hallmark EMP pathways were also differentially suppressed following immunotherapy in French bulldogs compared to other breeds, which mirrors MES-like de-differentiation in human GBM with acquired immunotherapy resistance [10,17,35].

Our study thus aims to bridge this gap to deepen the translational value of CG as a model of HG and reciprocally benefit canine patients. Using a combination of histology, immunohistochemistry, and tissue microarray (TMA)-based spatial transcriptomics, we investigated the dynamics between cell plasticity and immune TME in all three CG subtypes. We additionally emphasized correlations between histologic structures found in both high-grade CG and HG (microvascular proliferations and pseudopalisading necrosis), cancer cell phenotype, and immune cell polarization.

## Materials and Methods

### Sample selection

Thirty FFPE canine glioma samples were retrieved from the archives of the Cornell Animal Health Diagnostic Center (AHDC). Aside from grade (low-grade, high-grade), CG encompasses three subtypes defined by neoplastic cell morphology: astrocytoma, oligodendroglioma, and undefined glioma [34]. Because of the biological diversity of both CG and HG, and because of the canine propensity for developing high-grade oligodendrogliomas more than other grades and subtypes, cases were selected agnostic to patient signalment and tumor grade [30].

Ten astrocytomas (4/10 low-grade, 6/10 high-grade), ten oligodendrogliomas (1/10 low-grade, 9/10 high-grade), and ten undefined gliomas (3/10 low-grade, 7/10 high-grade) were selected blindly, agnostic to patient signalment and tumor grade. All cases were clinically evaluated and examined by a veterinary neurologist (ENG, JH, or GS), then submitted for histological evaluation, subtyping, and grading by a neuropathologist following the established high-grade/low-grade scheme (EAD/ADM) [34]. Hematoxylin-and-eosin (H&E)-stained slides associated with each case were retrieved and scanned at 20x for reviewing histomorphological features (VENTANA DP 200 slide scanner). Diagnoses and signalment are summarized in Table 1.

**Table 1:** Patient Signalment and Treatment History Correlated with T-and B-cell Immunohistochemistry Results.

| Subtype | Case number | Breed | Age | Sex | Grade | Survival | Treatment |  |  | T-cell IHC |  | B-cell IHC |  |
| --- | --- | --- | --- | --- | --- | --- | --- | --- | --- | --- | --- | --- | --- |
|  |  |  |  |  |  |  | High-dose steroids | Chemo-therapy | Radiation | CD3 | FOXP3 | PAX5 | CD20 |
| ASTROCYTOMA | Case 1 | Italian Greyhound | 10yo | FS | Low | 2 years | Y | N | N | (x) | (x) | (x) | (x) |
|  | Case 2 | Mixed Terrier | 12.5yo | FS | Low | 1 week | N | N | N | +/- | 0 | 0 | 0 |
|  | Case 3 | Boston Terrier | 7.5yo | MC | High | 3 weeks | Y | Y | N | + | 0 | +/- | +/- |
|  | Case 4 | American Bulldog | 11yo | FS | High | 1 week | Y | N | N | +/- | 0 | 0 | 0 |
|  | Case 5 | French Bulldog | 9yo | FS | High | 3 weeks | Y | N | N | +/- | 0 | 0 | 0 |
|  | Case 6 | Boston Terrier | 10.5yo | MC | High | 1 day | Y | N | N | 0 | 0 | 0 | 0 |
|  | Case 7 | Weimaraner | 3yo | FS | Low | 1 day | N | N | N | + | 0 | 0 | 0 |
|  | Case 8 | Mixed breed | 7yo | FS | High | 1 day | N | N | N | ++ | 0 | + | + |
|  | Case 9 | Mixed breed | 6yo | FS | High | NA | NA | NA | NA | +/- | +/- | 0 | 0 |
|  | Case 10 | Mixed breed | 7yo | MC | Low | 4 months | N | N | Y | + | +/- | 0 | 0 |
| OLIGODENDROGLIOMA | Case 11 | French Bulldog | 11yo | FS | High | 5 months | Y | N | N | 0 | 0 | 0 | 0 |
|  | Case 12 | Pitbull | 10yo | MC | Low | 2 days | N | N | N | + | 0 | 0 | 0 |
|  | Case 13 | Boston Terrier | 12yo | FS | High | 1 day | N | N | N | +/- | 0 | 0 | 0 |
|  | Case 14 | French Bulldog | 8yo | MC | High | NA | Y | N | N | +/- | +/- | 0 | 0 |
|  | Case 15 | Boston Terrier | 11yo | FS | High | 1 day | N | N | N | +/- | +/- | 0 | 0 |
|  | Case 16 | Boston Terrier | 8yo | NA | High | 2 months | Y | Y | N | +/- | 0 | 0 | 0 |
|  | Case 17 | Airedale Terrier | 7yo | FS | High | 3 weeks | N | N | N | +/- | +/- | +/- | +/- |
|  | Case 18 | Labrador Retriever | 8yo | MC | High | 1 day | N | N | N | + | 0 | 0 | 0 |
|  | Case 19 | Boxer | 10yo | MC | High | 1 day | N | N | N | +/- | 0 | 0 | 0 |
|  | Case 20 | French Bulldog | 4.5yo | FS | High | 2 days | Y | N | N | +/- | 0 | 0 | 0 |
| UNDEFINED GLIOMA | Case 21 | Golden Retriever | 1yo | MC | Low | 1 month | Y | N | N | (x) | (x) | (x) | (x) |
|  | Case 22 | Boston Terrier | 12yo | MC | High | NA | NA | NA | NA | +/- | 0 | 0 | 0 |
|  | Case 23 | English Bulldog | 14yo | FS | High | 3 days | Y | Y | N | 0 | 0 | 0 | 0 |
|  | Case 24 | Field Spaniel | 7yo | MC | High | 3 weeks | Y | Y | N | +/- | 0 | 0 | 0 |
|  | Case 25 | German Shepherd | 8yo | MC | High | 1 week | Y | N | N | +/- | 0 | 0 | 0 |
|  | Case 26 | French Bulldog | NA | MC | High | 3 days | Y | N | N | ++ | +/- | ++ | ++ |
|  | Case 27 | Labrador Retriever | 3mo | M | Low | 1 day | N | N | N | ++ | 0 | 0 | 0 |
|  | Case 28 | Australian Shepherd | 10yo | FS | High | 1 day | N | N | N | ++ | 0 | 0 | 0 |
|  | Case 29 | Dalmatian | 7yo | FS | High | 5 days | Y | Y | N | + | 0 | 0 | 0 |
|  | Case 30 | French Bulldog | 8.5yo | MC | High | 2 weeks | N | N | N | +/- | 0 | (x) | 0 |

Legend: NA: information not available, Y: yes, N: No, (x): no tumor tissue left, 0: absent, +/-: minimal to mild infiltrates, +: moderate infiltrates, ++: marked infiltrates

### Histomorphology-based categorization of neoplastic cells

To correlate cancer cell IHC labeling patterns and transcriptomics profiles with morphological features, neoplastic cells were subdivided into 4 categories using previously published criteria as a basis [18,34]: astrocyte-like cells (ALC), oligodendrocyte-like cells (OLC), stem-like cells (SLC), and undefined cells (UNC) (Supplemental Figure 1a). ALC were kite-shaped to spindle-shaped with moderate to abundant eosinophilic cytoplasm and a central, round to ovoid nucleus. OLC were spheroid with scant eosinophilic cytoplasm, a central hyperchromatic nucleus, and nested in clear spaces arranged in a honeycomb pattern. SLC were diminutive, ovoid to lacrimiform with minimal amounts of eosinophilic cytoplasm, and a central hyperchromatic nucleus. UNC had atypical features that did not fit any of the three categories above.

### Immunohistochemistry

The following antibodies were used for assessing immune cell infiltration: CD3 for T-cells (monoclonal antibody, clone LN10, Leica Microsystems, #PA0553-U), FoxP3 for regulatory T-cells (monoclonal antibody, clone FJK-16s, eBioscience, #14-5773-82), CD20 for B-cells (monoclonal antibody, clone IGEL/4524R, NeoBioTechnologies, #931-RBM14), PAX5 for B-cells (monoclonal antibody, clone 1EW, Leica Microsystems, #PA0552-U), MUM1/IRF4 for plasma cells (monoclonal antibody, clone EPR5653, Abcam, #ab124691), IBA1 for phagocytic cells and microglia (polyclonal antibody, FujiFilm Wako, #019-19741), and CD204 for anti-inflammatory phagocytes (monoclonal antibody, clone SRA-E5, Cosmo Bio, #KT022). For cell plasticity, the following were assessed: pan-cytokeratin (monoclonal antibody, clones AE1/AE3, DAKO, #M3515), E-cadherin (monoclonal antibody, clone 36, BD Biosciences, #610182), and vimentin (monoclonal antibody, clone Vim 3B4, DAKO, #M7020) as hallmark EMP markers. Finally, the following antibodies were used as CNS-specific markers of stemness and differentiation: OLIG2 (monoclonal antibody, clone EPR2673, ABCAM, #ab109186), GFAP (polyclonal antibody, DAKO, #Z0334), 2’,3’-cyclic-nucleotide 3’-phosphodiesterase (CNPAse; monoclonal antibody, clone 11-5B, Abcam, #ab6319), MAP2 (monoclonal antibody, Sigma-Aldrich, #45-M4403), SOX2 (monoclonal antibody, clone 20G5, Invitrogen, #MA1-014), and SOX10 (polyclonal antibody, Abcam, #ab180862). Immunohistochemistry (IHC) was performed using the Leica Bond Max Automated IHC Staining System, according to the manufacturer’s instructions (Leica Microsystems, Buffalo Grove, IL). Tissues were sectioned at a thickness of 5μm and deparaffinized with Bond Dewax Solution (Leica Biosystems, #AR9222). Detailed IHC protocols for most antibodies were previously described elsewhere; others (FoxP3, CD204, MUM1/IRF4) are validated in-house at the AHDC and routinely used for diagnostic purpose [44–49]. Summarized antibody specifications, reagents, and control tissues are available in Supplemental File S1.

FFPE sections were stained for each immune cell marker to assess their spatial distribution (T-cells, B-cells/plasma cells, IBA1^+^ and CD204^+^ phagocytes). SOX2 and SOX10 were also applied to whole sections to spatially correlate their expression with immune cell infiltration and cancer cell morphology. All other stemness and cell plasticity IHC antibodies (OLIG2, GFAP, CNPase, MAP2, E-cadherin, vimentin, pancytokeratin) were then applied to tissue microarrays, as described below.

### Tissue microarrays

Tissue microarrays (TMA) were built using a TMA molmed machine (3DHISTECH, Hungary). For whole-sections with tumor *in situ* (necropsy samples), regions of interest (ROI) were selected based on cancer cell morphology, SOX2 and SOX10 expression, and on immune cell infiltration assessed on H&E-and IHC-stained slides (CD3, CD20, PAX5, MUM1, IBA1, CD204). ROI were selected within each tumor and at the invasive front to ensure representation of intratumoral heterogeneity. For samples that consisted of tumor fragments (biopsy samples), ROI were selected based on the same criteria, though irrespective of their spatial tissue location (cases 1, 7, 9, 10; Table 1). One block was built per glioma subtype; each block consisted of 29 cores, for a total of 10 cases per block (Supplemental Figure 1b). Because TMA tissue cores can become heterogeneously exhausted depending on the thickness of the donor blocks, the total number of cores and cases available for evaluation per IHC marker is summarized in Supplemental File S1. One case consistently lacked sufficient tissue for proper evaluation thus was excluded from TMA evaluation (case 21; Table 1).

### Digital image analysis for standard histology slides (whole sections and TMAs)

The distribution of T-cells and phagocytes was analyzed in QuPath (v0.6.0 and 0.7.0) using the positive cell detection tool (CD3) and the thresholder (CD204, IBA1). When the thresholder was used, the following workflow and parameters were applied: (**i**) high resolution (1.86μm/px), channel: DAB, prefilter: Gaussian, smoothing sigma: 0.5-1, (**ii**) adjust detection threshold via live detection (0.20-0.25), (**iii**) create objects and classify objects as detection (new object type: detection, minimum object size: 10-15μm^2^, minimum hole size: 5μm^2^, split objects), (**iv**) manual quality check (QC) to reclassify inadequate detection (class: “Ignore”). For T-cell detection and quantification, a QC step was also performed first by verifying individual detection images (detection measurements) and reclassifying detections as needed (“Positive”/ “Negative”/“Ignore”), then cross-verified in-section by navigating the slide with the positive cell detection layer toggled on. For tumors that were well-delineated, the tumor itself and immediately adjacent neuroparenchyma were annotated as the region of interest. For tumors with a diffusely invasive pattern and biopsy samples, the whole section was annotated as the region of interest (Supplemental Figure 1c). Detections were used for generating heatmaps that reflect the distribution of each cell subtype (Supplemental Figure 1c). For density maps, the density radius was lowered to best represent the distribution of even sparsely scattered cells (50-125), transparency was set at 0.9, and the smoothness interpolation parameter was set as Bilinear. Semi-quantitative scores were used for evaluating SOX10 and SOX2 labeling intensity (0: absent, 1: minimal, 2: moderate, 3: marked). For either marker, the percentage of positive cells was determined by estimating the total tumor surface with labeling compared to the total tumor section area (0%: no labeling in any cell, 100%: positive labeling in all cells).

### Statistical analysis of immunohistochemistry results

T-tests were used to assess the statistical significance of scores and percentages between 2 groups (Student for presumed homoscedastic data, Welch for presumed heteroscedastic data). For comparison between more than 2 groups, a Kruskal-Wallis test (no distribution assumption) or one-way ANOVA test (Brown-Forsythe and Welch) was conducted. The median number of T-cells/mm^2^ across all subtypes (6.5 T-cells/mm^2^) was set as the cutoff value for categorizing tumors into T-cell^Hi^ and T-cell^Lo^ subgroups. The Mantel-Cox (log-rank) test was used to assess differences in OS between T-cell^Lo^ gliomas (< 6.5 T-cells/mm^2^) and T-cell^Hi^ gliomas (≥ 6.5 T-cells/mm^2^). OS was defined as the time elapsed between diagnosis and death or euthanasia. Direct correlation between OS and raw numbers of T-cells/mm^2^ was further assessed with a simple linear regression. A p-value of *p* ≤ .05 was considered statistically significant for all tests. The adjusted p-value was used instead of the raw p-value when available (*p*_adj_; Benjamini-Hochberg correction).

### TMA-based spatial transcriptomics (Xenium v1)

We designed a custom panel of 100 probes optimized for canine FFPE tissue targeting immune cell phenotypes and cancer cell plasticity and stemness (10x Genomics, Xenium V1). Target genes were selected based on previously published evidence of expression in canine glioma or other canine cancer(s), correlation between CG and HG, and EMP and immunosuppression in HG and in carcinomas [17,31,35]. Canine reference sets for annotations were GSE225599 and GSE252470, and the canine reference genome was ROS_Cfam_1.0 [50,51]. Gene signature groups included T-cell subtypes (*CD4*, *CD8*) and phenotypes (Th1, Th2 or Th17 response; T-cell exhaustion; effector/cytotoxic phenotype; memory/regulatory phenotype), B-cells, inflammatory receptors and chemokines (*JAK/STAT*), immunosuppressive macrophage phenotypes (*ARG1, SPP1*), glioma stemness/differentiation (*OLIG2, SOX10*), EMP (*ZEB1/2, TWIST1/2*), and malignancy signature genes divided into 4 subgroups: “upregulated in cancer” (*RAC1, EGFR*), “upregulated in French Bulldog glioma” (*PTX3, ADHFE1*), “upregulated in human glioma” (*ALDH1L2, SPARC*), “upregulated in both canine and human glioma” (*PDGFRA, SMARCA4*). The complete list of genes and subgroups defined for direct import and visualization in Xenium Explorer 4 is available in Supplemental File S2. Differentially expressed genes for each CG subtype are available in Supplemental File S3.

### Descriptive spatial transcriptomics data analysis

H&E images of each TMA were exported from QuPath-0.7.0 in OME-TIFF format and imported in the Xenium Explorer 4 software for direct overlap between histomorphological features and transcripts expression. For descriptive evaluation at the tissue core level, transcripts were visualized as either density maps, or points; the point size was kept consistent for each category across the three TMAs (“EMP”: size 11; “GStem”: size 12; “CD4”: size 15; “CD8”: size 14; “TAMs”: size 12). For descriptive evaluation at the cell level, we used differentially expressed gene (DEG) clusters generated by default Graph-Based clustering following Xenium v1 analysis: 16 DEG clusters for astrocytomas, 15 DEG clusters for oligodendrogliomas, and 22 DEG clusters for undefined gliomas. The raw DEG files were uploaded to Claude AI (Claude Fable 5) to retrieve statistically significant upregulated genes per cluster. Genes were sorted based on combined *p*_adj_ ≤ 0.05 and log2fold > 0, and each cluster was tentatively assigned a sub-category using a scoring system leveraging well-established gene signatures for EMP, glioma stemness, malignancy, T-cell exhaustion, Tregs, TAMs, and healthy glial cells [52–58]. Clusters and sub-categories underwent a manual quality check (QC) by a board-certified pathologist (SRN) and were given a definitive sub-category consistent with the most significantly upregulated genes per cluster, and according to combined histomorphological features on H&E, and most significantly upregulated genes per cluster (Supplemental File S2). Finally, we used *Smoothie*-based clustering (modules) for visually cross-verifying spatially correlated gene networks in relation to cell morphology and tissue architecture (Supplemental File S2).

### Xenium quality control

Per-cell QC metrics were computed with Scanpy (v1.10.1), and negative control probes, negative control codewords, and unassigned codewords were used to compute a noise ratio, defined as the sum of control and unassigned counts divided by total counts [59]. Cells were retained if they had at least 10 total transcripts, at least 5 detected genes, and a noise ratio of 0.01 or lower; control and unassigned features were then removed from the matrix, and QC metrics were recomputed on the filtered object. TMA cores were delineated manually by specifying the centroid of each core and assigning every cell to the core whose center lay within 1100 spatial units. Cells outside this radius of any core were discarded.

### Pseudobulk aggregation and differential expression

Pseudobulk profiles were generated from the filtered Xenium object by summing raw transcript counts across all cells within each TMA core. Cores containing fewer than 1000 cells were excluded. Aggregation was performed on TMAs annotated as tumor core, yielding 58 cores from 27 patients across the 100-gene panel, with a median of 11,340 cells per core. Core-level profiles were then summed within patients to give one profile per animal, ensuring statistical independence of replicates. The 27 cases included nine astrocytomas (4/9 low-grade, 5/9 high-grade; excluding case 6), eight oligodendrogliomas (1/8 low-grade, 7/8 high-grade; excluding cases 11 and 16), and ten undefined gliomas (2/10 low-grade, 8/10 high-grade).

Differential expression was performed with PyDESeq2 (v0.4.12) on patient-level integer counts using the design ∼condition with subtype as a three-level factor (astrocytoma as reference) and Cook’s distance-based outlier refitting enabled [60]. A single model was fitted across all 27 patients so that dispersions were estimated from the full dataset, and the three pairwise subtype contrasts were then extracted by Wald test. P-values were adjusted within each contrast by the Benjamini-Hochberg procedure and genes with adjusted *p* below 0.05 were called differentially expressed, giving fifteen genes for oligodendroglioma versus astrocytoma, seven for undefined versus astrocytoma, and one for undefined versus oligodendroglioma. For visualization, DESeq2 median-of-ratios normalized counts were log1p transformed and plotted per patient, with pairwise adjusted p-values annotated.

### Spatial gene module detection

Spatially covarying gene modules were identified with Smoothie [61]. Analysis was restricted to cells annotated as tumor core, and each TMA core was treated as an independent spatial dataset; cores with fewer than 1000 cells were excluded, leaving 58 cores with 1435 to 36,766 cells each: twenty-four astrocytoma cores (12/24 low-grade, 12/24 high-grade), thirteen oligodendroglioma cores (1/13 low-grade, 13/13 high-grade), and twenty-one undefined glioma cores (5/21 low-grade, 16/21 high-grade). Within each core, cells with fewer than 10 total transcripts were removed and genes with fewer than ten counts core-wide were dropped, retaining 75 to 100 panel genes per core. Counts were normalized to 1000 transcripts per cell and log1p transformed. Then the standard Smoothie pipeline was ran using a Gaussian kernel with standard deviation 40 μm and ‘min_spots_under_gaussian’ set as 25. A gene-gene network was built by retaining edges with a Pearson coefficient above 0.3 and clustering power set as four. Modules were determined based on gene network clustering.

### Module scoring and downstream comparisons

Spatial gene modules with two or more genes were taken from the Smoothie network. This resulted in 13 modules that were given descriptive labels (G-MAL, TAM-LEX, CTL-NK, iMES-EX, OPC-like, TAN-TRF, iTR-MES, MES-IL6, EPC-BR, AC-RA, SNAI1-IS, CHK-T, and APM). Counts were normalized to 1000 transcripts per cell and log1p transformed. Each module was then scored in every cell as the mean of its member genes’ z-scored expression, with gene means and standard deviations computed once across all cells and z-scores clipped at plus or minus 10.

To avoid pseudo-replication, cell-level scores were aggregated in two steps. Cores with fewer than 1000 cells and cells from patients lacking subtype annotation were excluded; remaining cells were averaged within each TMA core, and within each core-region combination in parallel, retaining the fraction of cells with a positive score alongside the mean. Core-level means were then averaged up to one value per patient, weighing each core equally regardless of cell count. Subtype comparisons used the patient-level table restricted to tumor core tissue; region comparisons used the core-level, region-stratified table within each subtype.

For each module, groups were compared by Kruskal-Wallis test followed by two-sided pairwise Mann-Whitney U tests, with p values Bonferroni corrected across the pairwise contrasts within a module and additionally Benjamini-Hochberg corrected across all module-by-contrast tests in a given table; significance was set at 0.05. Group summaries were displayed as heatmaps of module means z-scored within each module across groups. Module co-occurrence was assessed by Spearman correlation between all module pairs across patients, with Benjamini-Hochberg correction across pairs. The TAM-LEX module was additionally correlated against every other module across patients (Spearman, Benjamini-Hochberg corrected) and plotted against the iMES-EX, TAN-TRF, and iTR-MES modules with a least-squares fit, coloring patients by subtype.

## Results

### Canine glioma displays subtype-specific preferential differentiation into defined plastic cell phenotypes, and differential expression of stemness marker SOX10

First, we investigated the expression of two hallmark markers of human glioma stemness: SOX2 and SOX10 [62,63]. SOX10 expression has been previously observed in CG in correlation with other markers of oligodendrocyte precursors (OLIG2, NG2) [47,64]. Importantly, SOX10 has been repeatedly described as the gatekeeper of cell plasticity and stemness in humans, and is thought to be useful in combination with OLIG2 for prognostic purposes [62,65,66].

Whole-section IHC analysis showed that SOX2 was expressed consistently across all three tumor types, most often diffusely, and at a moderate to strong labeling intensity (Fig. 1a). SOX10, on the other hand, showed a higher percentage of expression in oligodendrogliomas and in undefined gliomas, with an average moderate labeling intensity (Fig. 1b). Canine breeds predisposed to glioma (American bulldogs, French bulldogs, Boston terriers, Boxers) exhibited neither a higher percentage of SOX10-positive cells, nor increased labeling intensity (Fig. 1b). SOX10 labeled SLC and UNC most frequently, though well-differentiated OLC expressed SOX10 as well (Fig. 1c). In tumors with heterogeneous labeling, SOX10 was preferentially expressed strongly at the invasive front of tumors, and in pseudo-palisading neoplastic cells (Supplemental Figure 2a). Microvascular proliferations (MP) had SOX2 nuclear labeling but lacked SOX10 labeling (Supplemental Figure 2b).

**Figure 1.**
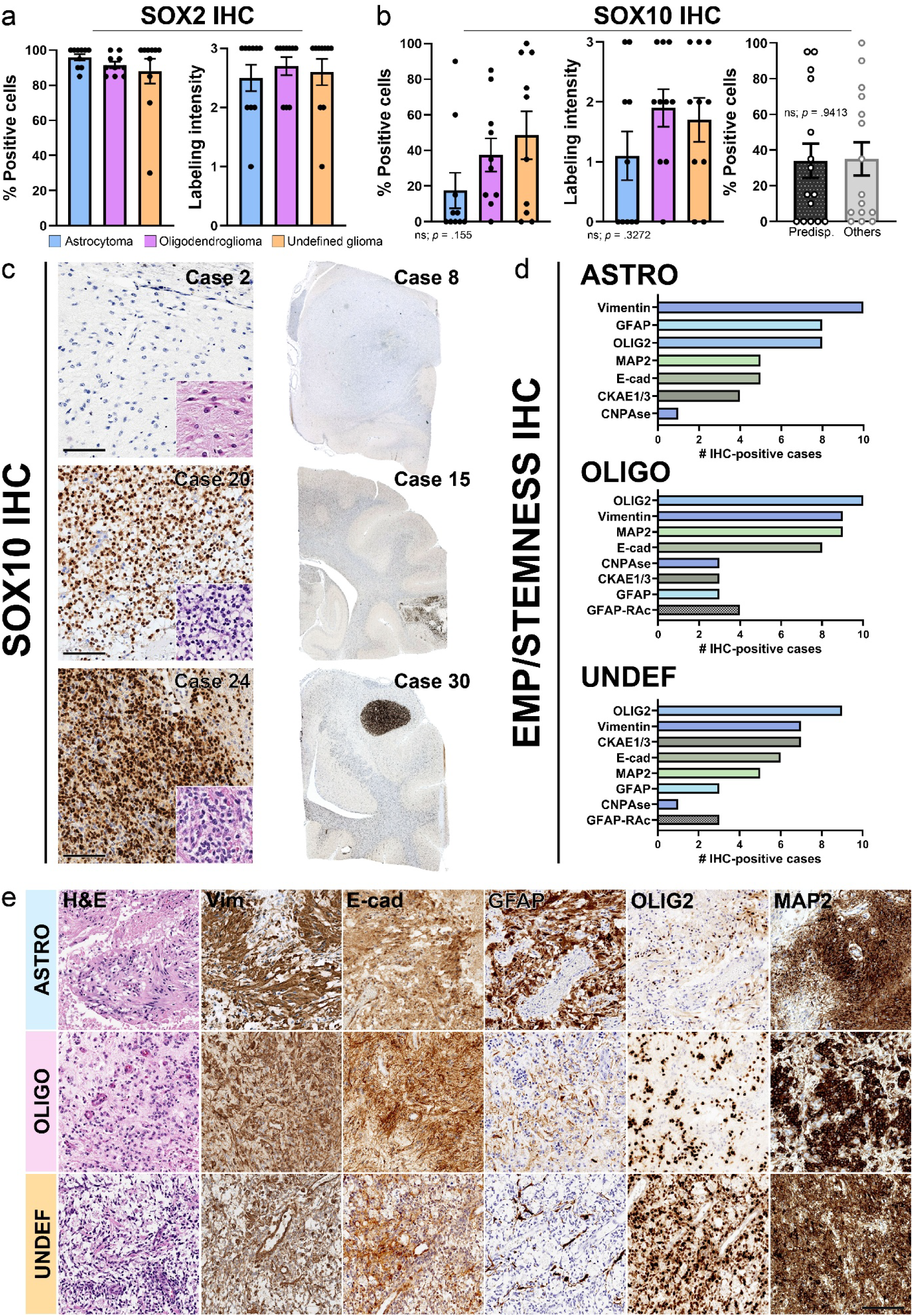
Immunohistochemistry-based assessment of glioma stemness and cell plasticity markers. **(a)** The percentage of SOX2^+^ cells and labeling intensity were both high across all gliomas, regardless of subtype and grade. **(b)** The percentage of SOX10^+^ cells was highest in undefined gliomas. The intensity of labeling was lowest in astrocytomas, and highest in oligodendrogliomas and undefined gliomas. No defined IHC labeling trend emerged between predisposed breeds and others. **(c)** Astrocytomas frequently had sparse to absent SOX10 labeling. Oligodendrogliomas often had heterogeneous SOX10 labeling; well-differentiated OLC commonly had mixed moderate to strong SOX10 nuclear labeling. In undefined gliomas, the percentage of SOX10^+^ neoplastic cells was often >90%. Undefined gliomas often consisted of SLC and UNC that resembled neither astrocytes, nor oligodendrocytes. **(d)** IHC results per glioma subtype. GFAP labeled reactive astrocytes (RAc)-like cells frequently in both oligodendrogliomas (OLIGO) and undefined gliomas (UNDEF). **(e)** Illustration of IHC-based cell phenotyping for each CG subtype. Astrocytomas (ASTRO) typically had a GFAP^+^Vim^+^E-cad(cytoplasmic)^+^ phenotype most frequently seen in UNC and ALC, while oligodendrogliomas and undefined gliomas most frequently were OLIG2^+^SOX10^+^MAP2^+^, with vimentin labeling in UNC cells surrounding areas of geographic necrosis. This suggested a more-mesenchymal phenotype in astrocytomas, and a more stem-like phenotype in oligodendrogliomas and undefined gliomas. Scale bar: 100μm.

We evaluated neoplastic cell phenotype further by conducting IHC for E-cadherin, vimentin, pancytokeratin, GFAP, OLIG2, MAP2, and CNPase applied to CG TMA (Fig. 1d). Astrocytomas most strongly expressed GFAP, vimentin, and OLIG2, followed by MAP2, E-cadherin, and pancytokeratin (Fig. 1d). Oligodendrogliomas highly expressed OLIG2, vimentin, MAP2, and E-cadherin, and less frequently labeled with pancytokeratin, GFAP, and CNPase (Fig. 1d). Undefined gliomas similarly had high expression of OLIG2, vimentin, and pancytokeratin, as well as E-cadherin and MAP2; GFAP and CNPase were lowly expressed (Fig. 1d).

For cell plasticity markers (E-cadherin, vimentin, pancytokeratin), labeling patterns were then correlated with cell morphology for each CG subtype, with an emphasis on the sub-cellular localization of E-cadherin. Pancytokeratin and E-cadherin are hallmarks of EMP, and labeling has previously been described in human GBM [67,68]. E-cadherin labeling was often cytoplasmic rather than membranous, a feature described in de-differentiated cells that reside in a “hybrid” epithelial-mesenchymal (E/M) phenotype (Fig. 1e) [69]. In astrocytomas, pseudopalisading UNC cells delineating areas of necrosis commonly had cytoplasmic and membranous E-cadherin labeling (Fig. 1e). Membranous and/or cytoplasmic E-cadherin labeling was also detected in UNC surrounding sprouting vascular structures in all CG subtypes. No cross-labeling with GFAP was observed (Supplemental Figure 2c).

Oligodendrogliomas generally lacked E-cadherin labeling in OLC, SLC, and UNC; when detected, E-cadherin labeled perivascular UNC (Fig. 1e, Supplemental Figure 2c). In undefined gliomas, the E-cadherin labeling pattern was consistently both cytoplasmic and membranous, with moderate labeling intensity (Fig. 1e).

Vimentin was most highly expressed by UNC and ALC in astrocytomas (Fig. 1e). In oligodendrogliomas and undefined gliomas, vimentin most frequently labeled SLC and UNC. Though the percentage of positive neoplastic cells varied, vimentin expression was typically strong in pseudopalisading SLC and UNC cells, following the same pattern as cytoplasmic E-cadherin and SOX10 (Fig. 1e, Supplemental Figure 2d). The exception was well-differentiated OLC, which lacked vimentin labeling in all tumor subtypes (Supplemental Figure 2d). No overt breed predisposition was observed, and no difference in labeling pattern and intensity was seen between the center of the tumor and the invasive front in any CG subtype. Finally, though we noted moderate to strong cytoplasmic pancytokeratin labeling in undefined gliomas more frequently than other CG subtypes, we found that the labeling pattern strongly mirrored this of GFAP and often mapped to cells with a reactive astrocyte morphology. Combined with aberrant pancytokeratin labeling in the glia limitans of a healthy control canine brain, we interpreted pancytokeratin labeling in CG as cross-labeling in our cohort (Supplemental Figure 2e).

For standard CG diagnostic markers (GFAP, OLIG2) and MAP2, IHC labeling patterns have been extensively characterized previously, and our cohort followed these known trends [34,44]. With regard specifically to cell morphology, MAP2 labeled SLC and UNC in all CG subtypes (Fig. 1d). GFAP expression was expectedly highest in astrocytomas; in oligodendrogliomas and undefined gliomas, GFAP often highlighted cells that resembled reactive astrocytes rather than neoplastic ALC (Fig. 1e). These reactive astrocytes (“RAc”) had a standard astrocytic morphology with slender cytoplasmic processes wrapping around neoplastic cells (Fig. 1d). Though OLIG2 labeling was not found more frequently in oligodendrogliomas or undefined gliomas than in astrocytomas, the labeling intensity and percentage of labeled cells were higher in oligodendrogliomas and undefined gliomas (Supplemental Figure 2f), and French Bulldogs consistently had strong and extensive OLIG2 labeling (Supplemental File S1). CNPase IHC was broadly negative; in positive cases, labeling was extremely sparse, faint, and limited to discrete clusters of SLC. The neuroparenchyma in “healthy” control TMA cores had adequate labeling for all markers.

In summary, we identified cross-CG subtype and within-patient heterogeneous phenotypic niches. Astrocytomas had a marked GFAP^+^Vim^+^E-cad(cytoplasmic)^+^ phenotype with variable SOX10 labeling, that typically highlighted ALC or UNC. Oligodendrogliomas and undefined gliomas mostly consisted of OLIG2^+^SOX10^+^MAP2^+^ cells, with vimentin labeling highlighting UNC in areas of pseudopalisading necrosis, as well as frequent occurrence of UNC cells with cytoplasmic E-cadherin labeling surrounding *de novo* formation of MP. Astrocytomas predominantly exhibited a trend toward acquisition of a more-mesenchymal phenotype, while oligodendrogliomas and undefined gliomas preferentially maintained a stem-like state, with OLC more frequently occurring in oligodendrogliomas and SLC in undefined gliomas. Therefore, IHC evaluation demonstrated the existence of heterogeneous niches across subtypes and across patients, as found in HG.

### Canine glioma exhibits heterogeneous, subtype-specific more-mesenchymal and more stem-like neoplastic niches at the transcriptional level

To deepen our assessment of neoplastic cell phenotypes and segregation into niches assessed by IHC, we used Graph-Based (GB) gene clustering to correlate upregulated transcripts with atypical histomorphological features and IHC protein expression profiles. GB clusters that both had a dominant cell plasticity (EMP) or glioma stemness (GStem) component and were expressed by cancer cells in Xenium Explorer 4 were considered relevant (Supplemental File S2).

Astrocytomas yielded two GStem clusters (clusters 5 and 8) and two EMP clusters (clusters 3 and 4) (Fig. 2a). Three out of 9 evaluable cases upregulated GStem clusters extensively: case 8 (cluster 8), and cases 9-10 (cluster 5). SLC preferentially upregulated cluster 8, while ALC upregulated cluster 5 (Fig. 2b). Inter-patient variability was observed for all 4 clusters: EMP cluster 3 was most widely upregulated in case 10; EMP cluster 4 was most strongly upregulated in cases 4 and 5; GStem cluster 5 was strongly upregulated in case 9; and GStem cluster 8 was strongly upregulated in case 8. Cases 2, 4, and 5 upregulated EMP clusters more extensively than GStem clusters. EMP cluster 3 specifically mapped to proliferative intratumoral vessels (Fig. 2c). Cores that expressed GStem clusters 5 and/or 8 generally upregulated EMP clusters 3 and/or 4. In case 8, EMP cluster 4 was preferentially upregulated by intratumoral cores, while GStem cluster 5 was preferentially upregulated at the invasive front (Fig. 2d). In other cases, tumor cores selected from the tumor center and invasive front equivalently upregulated GStem and EMP clusters.

**Figure 2.**
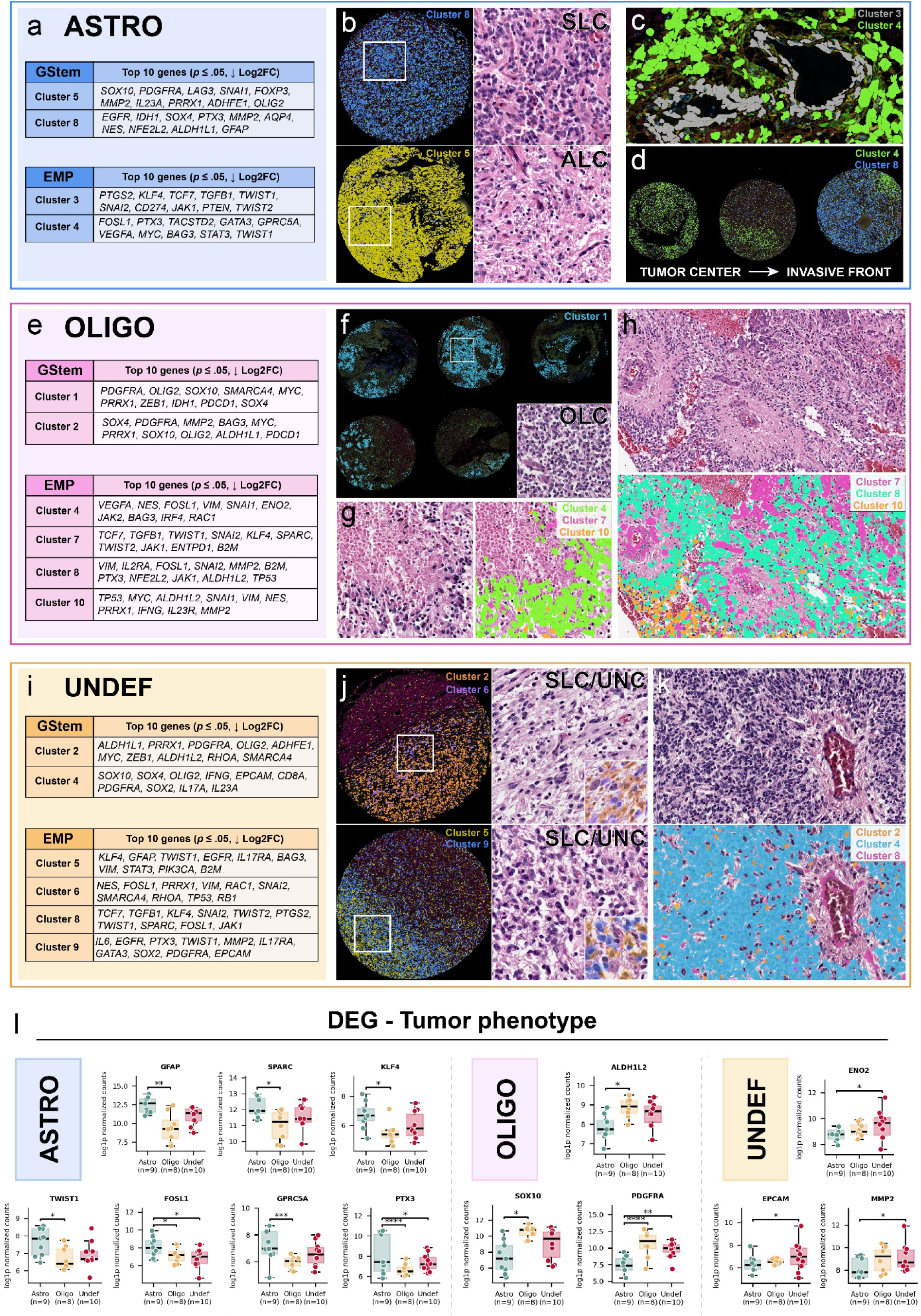
Subtype-specific graph-based clustering gene signatures correlated with histomorphological features and differentially expressed genes across subtypes. **(a-d) Astrocytomas.** Astrocytomas upregulated two stem-like (GStem) and cell plasticity-like (EMP) clusters. **(b)** These clusters were differentially expressed across patients and were upregulated as much by ALC as by SLC. **(c)** EMP cluster 4 mapped most frequently to neoplastic cells, while EMP cluster 3 distinctly highlighted proliferative intratumoral vessels. **(d)** Clusters 4 and 8 displayed a gradient of differential expression between tumor center and invasive front. **(e-h) Oligodendrogliomas**. **(e)** GStem clusters distinctly included PDGFRA, OLIG2, and SOX10, compatible with an OPC-like phenotype. Increased diversity of EMP phenotypes was appreciated, with 4 distinct GB clusters. **(f)** The extent of cluster 1 expression varied across cases; cluster 1 was generally upregulated by OLC. **(g-h)** EMP clusters were differentially expressed within one tumor and in specific areas (pseudopalisading necrosis) and predominantly mapped to SLC and UNC. EMP cluster 7 highlighted proliferative intratumoral vessels and MPs. **(i-k) Undefined gliomas**. **(i)** Similarly to oligodendrogliomas, undefined gliomas had 4 EMP-like signatures. **(j)** Heterogeneous GStem and EMP niches co-localized differently within one tumor and across cases. **(k)** GStem clusters generally mapped to SLC. **(l)** Differentially expressed genes (DEG) across CG subtypes. *: *p* = ≤ .05, **: *p* = ≤ .01, ***: *p* = ≤ .001, ****: *p* = ≤ .0001 (Benjamini-Hochberg).

In oligodendrogliomas, we identified two GStem clusters (clusters 1 and 2), and four EMP clusters (clusters 4, 7, 8, and 10) (Fig. 2e). Oligodendrogliomas had more frequent and extensive upregulation of GStem clusters than astrocytomas. Though GStem cluster 1 was expressed by OLC and SLC in almost all cases (cases 11, 13, 14, 15, 16, 17, 19, 20), upregulation was most extensive in cases 13, 14, 15, and 19 (Fig. 2f). Cluster 2, on the other hand, was strongly associated with OLC in case 20, and more sparsely upregulated by OLC and SLC in other cases. EMP cluster 4 was highly expressed in cases 13-15 and mapped to neoplastic cells surrounding necrotic areas, arranged or not into pseudopalisades (Fig. 2g). EMP cluster 7 mapped to MP and to intratumoral blood vessels (Fig. 2h). EMP cluster 10 was highly specific to non-palisading aggregates of SLC and UNC in case 17 (Fig. 2h). EMP cluster 8 was a small 11-gene cluster specifically expressed by UNC in a single tumor core (case 17), and by pseudopalisading UNC in another tumor core (case 18) (Fig. 2h). SLC and OLC expressing clusters 8 and 10 were also found sparsely scattered throughout most other cases, often in association with areas of edema, necrosis, and proliferative intratumoral vessels. Case 14 upregulated all EMP clusters (clusters 4, 8, 10) and GStem gene clusters concurrently (clusters 1 and 2). *OLIG2* was generally expressed by cancer cells upregulating GStem and EMP clusters. IHC-based OLIG2 protein expression strongly correlated with overexpression of GStem clusters 1 and 2. One case (case 15) additionally overexpressed a neoplastic astrocyte-like cluster (cluster 6) though *OLIG2* was consistently expressed by cancer cells (Supplemental File S2).

Finally, undefined gliomas yielded two GStem clusters (clusters 2 and 4), and four EMP clusters (clusters 5, 6, 8, 9) (Fig. 2i). Undefined gliomas had the highest degree of inter-patient variation regarding EMP clusters upregulation. GStem cluster 2 was upregulated by cases 22 and 24, and GStem cluster 4 was upregulated by cases 21 and 30. EMP cluster 5 was upregulated by case 26 only, concurrently with EMP cluster 9 (Fig. 2j). Cases 22 and 24 upregulated EMP cluster 6 concurrently with GStem cluster 2; in either case, cluster 6 was more extensively expressed at/near the invasive front, and more sparsely expressed by scattered neoplastic cells in other cores (Fig. 2j). GStem and EMP clusters were upregulated by both SLC and UNC. Finally, EMP cluster 8 mapped to both intratumoral vessels and vessels at the invasive front in all cases, suggesting ties to intratumoral vascular proliferation (Fig. 2k). Therefore, GB-clustering analysis supports that although EMP and GStem gene clusters slightly differ across CG subtypes, astrocytomas extensively upregulate more-mesenchymal EMP gene signatures, and oligodendrogliomas upregulate GStem signatures more frequently. Undefined gliomas specifically yielded a more heterogeneous blend of EMP and GStem gene clusters. Each CG subtype additionally had one EMP-like signature that mapped to proliferating intratumoral vessels and, when present in TMA cores, to MP.

These findings were reinforced by cross-CG subtype DEG analysis (Fig. 2l), which revealed significant upregulation of the major glioma stemness-associated gene *PDGFRA* in both oligodendrogliomas and undefined gliomas compared to astrocytomas. Astrocytomas stood out again by differential upregulation of EMP genes not only indicating residence in a more-mesenchymal state (*TWIST1, PTX3, FOSL1*) and confirming an astrocytic signature (*GFAP, SPARC*), but also a specific Yamanaka factor (*KLF4*), and increased propensity for invasion (*GPRC5A*). DEG also confirmed differential upregulation of *SOX10* in oligodendrogliomas and undefined gliomas, as quantified in tissue IHC analysis. Additionally, oligodendrogliomas upregulated *SOX4*, which is typically associated with neural progenitor cell (NPC)-like populations in human GBM [18]. Undefined gliomas uniquely upregulated *MMP2, ENO2, EPCAM*, and *IDH1*, suggesting that although oligodendrogliomas and undefined gliomas broadly reside in a more stem-like state, their transcriptional phenotype may differ. Thus, each CG subtype showed inter-patient diversity regarding cell plasticity and glioma stemness, together with within-patient heterogeneous intratumoral niches. Subtype-specific phenotypic trends and intratumoral niche heterogeneity within CG subtype, across patients, and within one tumor emerged at the transcriptional level.

### All canine glioma subtypes host exhausted T-cells and TAMs that often map to EMP-heavy niches

With few exceptions (cases 3, 8, 9, 19, 17, and 26), intratumoral CD3^+^ T-cells were sparse and randomly scattered, correlating with established CG literature [43]. Astrocytomas exhibited a strong though non-statistically significant trend toward higher numbers of infiltrating T-cells/mm² than oligodendrogliomas and undefined gliomas (Brown-Forsythe test, *p* = .1166) (Fig. 3a). In all subtypes, dogs that had received treatment showed a trend to decreased intratumoral T-cell infiltration, though not statistically significant either (Welch’s t-test, *p* = .2022) (Fig. 3a). Intratumoral T-cell infiltration rates did not correlate with OS based on a Mantel-Cox test (*p* = .7274) and simple linear regression (*p* = .8609) (Fig. 3b).

**Figure 3.**
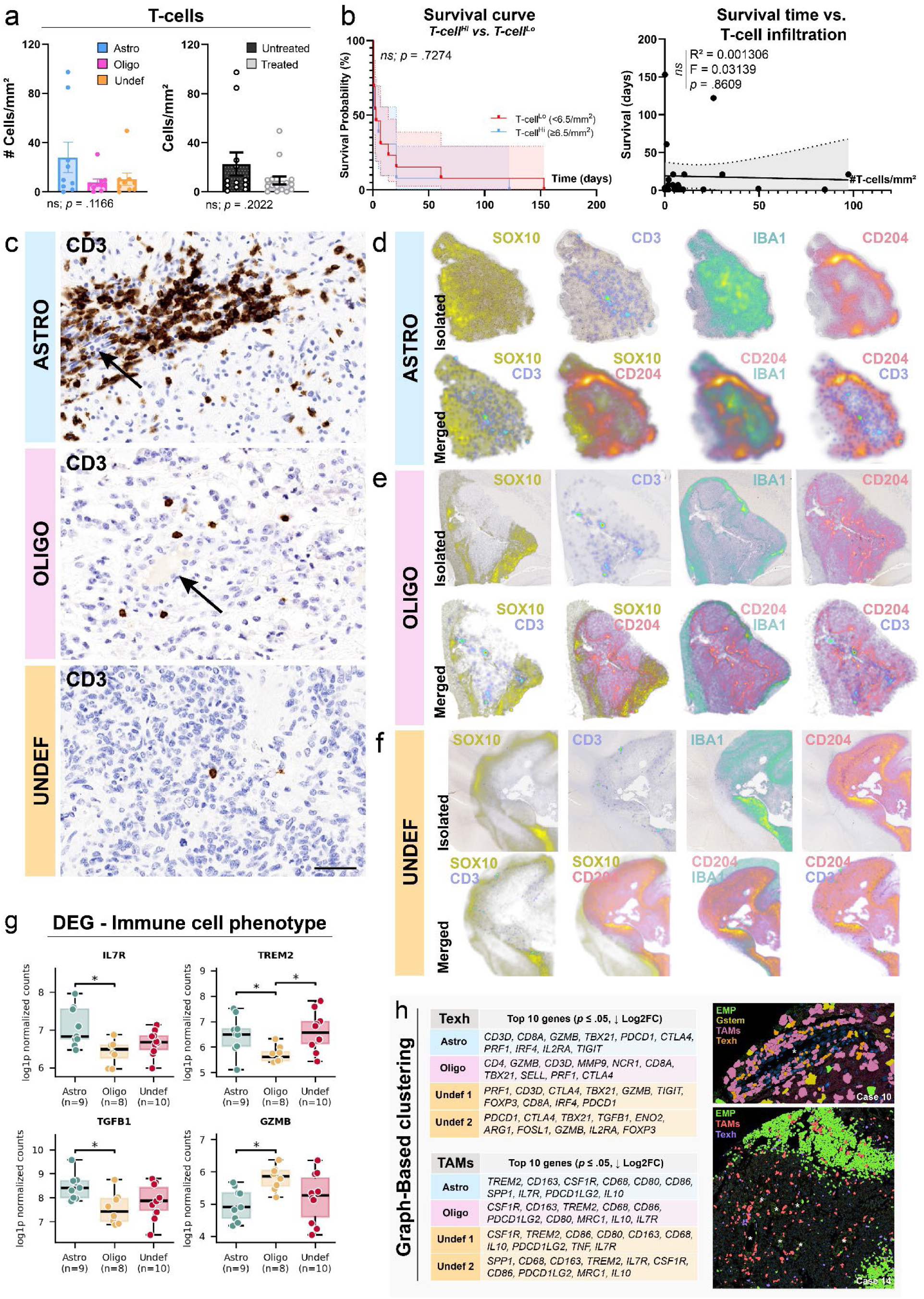
Immune microenvironment across CG subtypes. **(a-c) T-cell infiltration in relation to CG subtype and survival**. **(a)** Though non-statistically significant, astrocytomas showed a strong trend to increased T-cell infiltration compared to other subtypes. A moderate trend to increased intra-tumoral T-cell numbers was seen in untreated dogs compared to treated dogs. **(b)** Intra-tumoral T-cell infiltration did not correlate with survival time either based on infiltration density (< 6.5 cells/mm^2^, ≥ 6.5 cells/mm^2^) or raw numbers of T-cells/mm^2^. **(c)** CD3^+^ T-cells were more numerous in astrocytomas than in other subtypes and often perivascular (ASTRO). Intratumoral T-cells were less abundant in oligodendrogliomas and either randomly scattered or aggregated in areas of necrosis and around vessels (OLIGO). Undefined gliomas were sparsely infiltrated by T-cells (UNDEF). Black arrows: intratumoral blood vessels. **(d-f) Spatial distribution of T-cells (CD3), immunosuppressive macrophages (CD204), and macrophages and activated microglia (IBA1) in relation to SOX10 expression.** Heatmaps highlighted T-cell infiltration hotspots within astrocytomas **(d)** more frequently than in oligodendrogliomas **(e)** and undefined gliomas **(f)**. In all CG subtypes, CD204^+^ cells widely infiltrated tumors and often aggregated near central areas of necrosis. IBA1^+^ cells tended to infiltrate CD204^+^ cell-poor regions and reciprocally for CD204^+^ cells in IBA1^+^ cell-poor regions. SOX10 was generally expressed at the invasive front of tumors with heterogeneous CD3^+^ T-cells were frequently excluded from SOX10-positive areas. **(g-h) Cross-subtype immune cell phenotypes. (g)** Astrocytomas upregulated *TGFB1* and *IL7R*; oligodendrogliomas upregulated *GZMB*; and undefined gliomas and astrocytomas upregulated *TREM2*. **(h)** Each CG subtype upregulated exhausted and/or dysregulated T-cell (Texh) and immunosuppressive macrophages (TAMs) gene signatures. TAMs surrounded intratumoral blood vessels and often clustered with exhausted T-cells and regulatory T-cells in all subtypes (case 10). TAMs additionally mapped frequently to EMP-heavy niches, which occurred heterogeneously within one tumor and across cases (case 14). *: blood vessels.

In astrocytomas, CD3^+^ T-cells often formed peri-vascular cuffs (PVC) and were frequently found in dense aggregates that obscured degenerate neoplastic cells (Fig. 3c, 3d). T-cells were sparser in oligodendrogliomas, though also coalesced into small lymphoid clusters that followed a similar pattern (Fig. 3c, 3e). Undefined gliomas had low numbers of intratumoral T-cells that were often sparse and isolated, and tended to be confined to the nearby meninges and/or to the invasive front (Fig. 3c, 3f).

Histologically, there was no consistent evidence of overlap or exclusion of CD3^+^ T-cells by CD204^+^ cell-rich areas (Fig. 3d-f). Though CD3^+^ T-cells often accumulated preferentially within areas with mild to absent SOX10 labeling (Fig. 3d-f), no positive or negative correlate was found between %SOX10 labeling and CD3^+^ T-cell numbers either (Supplemental Figure 2g). CD204^+^ cells widely infiltrated all tumors; tumors in whole-section samples were often surrounded by IBA1^+^ cells (Fig. 3d). IBA1^+^ cells tended to fill CD204^+^ cell-poor areas and reciprocally in an interlocking fashion (Fig. 3d-f). CD204^+^ additionally displayed preferential accumulation not only within areas of necrosis, but also near MPs (Supplemental Figure 2h). FoxP3^+^ cells were rarely seen and in low numbers when present (< 1 cell in a digital 2.37mm^2^ field). Only 6 dogs had FoxP3^+^ cells, with a slight bias toward oligodendrogliomas (cases 9, 10, 14, 15, 17, and 26; Table 1). B-cells were generally absent; 4 dogs had CD20-and PAX5-positive B-cells, which mapped to PVC (cases 3, 8, 17, 26; Table 1).

Cross-CG subtype DEG analysis additionally highlighted three major immune-adjacent factors with differential upregulation between astrocytomas and oligodendrogliomas (Fig. 3g): *IL7R*, whose upregulation in human GBM is associated with decreased OS, was upregulated in astrocytomas [70]. *TGFB1* was upregulated in astrocytomas as well, correlating with its subtype-specific tendency to host more-mesenchymal niches. *GZMB,* on the other hand, was upregulated by oligodendrogliomas, suggesting higher infiltration rates by cytotoxic effectors T-cells. *TREM2*, which is upregulated by some TAMs and by myeloid-derived suppressor cells (MDSCs), was differentially upregulated in both astrocytomas and undefined gliomas compared to oligodendrogliomas. This supports either the existence of a denser TAM population in the former two, or the predominance of a phenotypically distinct TAM population in the latter. GB clustering analysis solidified these trends by highlighting distinct regulatory T-cell/T-cell exhaustion (“Texh”) and immunosuppressive macrophages (“TAM”) signatures in all three CG subtypes (Fig. 3h). Inter-patient variability and intratumoral heterogeneity were observed when overlapping GB clusters to TMA cores, reinforcing trends identified with EMP and GStem GB clusters. In all CG subtypes, TAMs were found in perivascular niches and near areas of necrosis, mirroring the CD204 IHC findings (Fig. 3h). In cores with dual EMP and glioma stemness clusters upregulation, TAMs often showed preferential accumulation in areas with strong EMP activation and spatially co-occurred with clusters of exhausted T-cells and/or Tregs (Fig. 3h).

### EMP and cancer cell stemness spatially correlate with T-cell exhaustion and immunosuppressive macrophages infiltration at the transcriptional level

At the core level, we descriptively correlated canonical transcripts for EMP (*ZEB1/2, TWIST1/2, SNAI1/2, EPCAM, TGFB1*), glioma stemness (*SOX10, OLIG2),* CD4 T-cells (*CD3D, CD4*), exhausted CD8 T-cells (*CD3D, CD8A, CD244, LAG3, PDCD1, CTLA4, HAVCR2, TIGIT*), and TAMs (*CD68, CD80/86, ARG1, CD163, CSF1R, MRC1, NFE2L2, TREM2)*. Astrocytomas had moderate EMP upregulation in 3 cases (3, 4, 5) and extensive EMP upregulation in 4 cases (7, 8, 9, 10) (Supplemental Figure 3a). *SOX10* was proportionally upregulated moderately to extensively in cases 5, 9, and 10; *OLIG2*, however, was consistently upregulated in cores with a strong EMP signature in all CG subtypes. In all CG subtypes, EMP transcripts upregulation also correlated with infiltration by exhausted CD8 T-cells, CD4 T-cells, and TAMs. Cases 3, 4, 5, 7, 8, 9, and 10 were most representative of this trend. Oligodendrogliomas and undefined gliomas showed similar correlations, though with some subtype-specific variation (Supplemental Figure 3b-c). Both subtypes more consistently upregulated EMP transcripts moderately to extensively (cases 13, 14, 15, 17, 18, 19, 20, 21, 22, 24, 26, 27, 29, 30). Oligodendrogliomas consistently expressed *SOX10* strongly in all tumor cores.

### More-mesenchymal tumor phenotypes and immunosuppression co-occur in heterogeneous niches across all CG subtypes

Because co-occurrence of defined cell types and GB clustering described in tissue sections are not sufficient to surmise significant correlations between T-cell exhaustion, Tregs, TAMs, and EMP/glioma stemness, we expanded our analysis by leveraging *Smoothie*-based transcriptomics evaluation [61]. *Smoothie* complements our descriptive analysis by detecting networks of concurrently upregulated genes in a cell type-agnostic fashion, allowing for identifying mixed intratumoral niches. Focusing on tumor cores only, we identified 13 distinct modules often combining genes derived from EMP and/or glioma stemness, and immune cell identity and polarization (Fig. 4a, Supplemental File S2).

**Figure 4.**
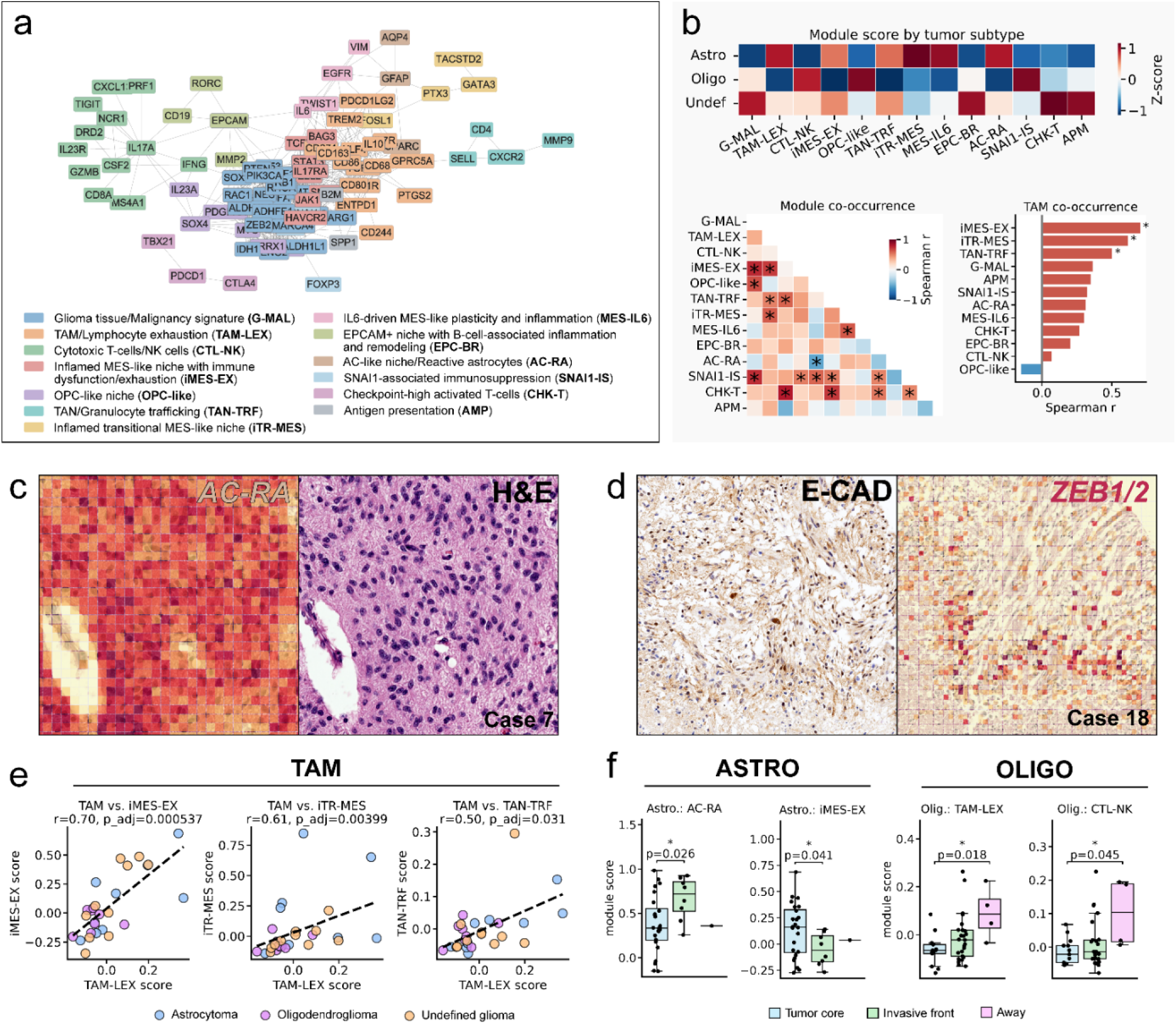
Module-level gene expression analysis. **(a)** *Smoothie*-based analysis yielded 13 distinct modules that frequently combined cell plasticity and stemness features with immune dysfunction and immunosuppression. **(b)** Some modules displayed strong ties to one CG subtype. TAM-LEX, iTR-MES, MES-IL6, AC-RA for astrocytoma; CTL-NK, OPC-like, and SNAI1-IS for oligodendroglioma; and G-MAL, EPC-BR, CHK-T, and APM for undefined glioma. **(c)** ALC in astrocytomas typically upregulated the AC-RA module. Heatmap, 10×10μm bins. **(d)** Neoplastic cells with cytoplasmic E-cadherin expression upregulated ZEB1/2. Heatmap, 10×10μm bins. **(e)** Per-case detail of TAM module co-occurrence with iMES-EX, iTR-MES, and TAN-TRF. Oligodendroglioma cases cluster near the origin of either axis, supporting a true invert TAM/oligodendroglioma and TAM/OPC-like relation. Astrocytomas exhibit more heterogeneous trends than undefined gliomas. **(f)** In astrocytomas, the AC-RA module is upregulated more frequently at the invasive tumor front, and the iMES-EX module is preferentially upregulated within tumors. Oligodendrogliomas show a trend to sequestering TAM-LEX and CTL-NK populations at the periphery of the tumor.

We first investigated the degree of correlation between CG subtype and *Smoothie* modules (Fig. 4b). Astrocytomas were strongly associated with the following modules: TAM-LEX, iTR-MES, MES-IL6, and AC-RA (Fig. 4b). The TAM-LEX module combined a standard TAM signature (*CSF1R, CD68, CD163, CD80, CD86, TREM2, KLF4, GPRC5A*), TAM-derived immunosuppressive genes (*TGFB1, IL10, PTGS2, CD274, PDCD1LG2*), and T-/NK-cell exhaustion markers (*CD244, ENTPD1, IL17R*). MES-IL6 was a small four-gene EMP-adjacent module that combined *EGFR, IL6, VIM*, and *TWIST1*, which indicated more-mesenchymal differentiation potentially driven by IL-6 [71]. The AC-RA module (*GFAP, AQP4, SPARC*) was the most widely upregulated across all astrocytoma cores and was expressed mainly by ALC (Fig. 4c). Oligodendrogliomas showed significant association with 3 modules: CTL-NK, OPC-like, and SNAI1-IS. The OPC-like module consisted of genes canonically expressed by oligodendrocyte progenitor cells (*OLIG2, SOX10, PDGFRA*), and a combination of EMP, stemness, and malignancy genes (*PRRX1*, *SOX4, MYC, IL23A*) [18]. The CTL-NK module consisted of genes indicating a mix of activated, likely exhausted CD8 T-cells; B-cells; and NK cells that spatially co-occurred with upregulation of *DRD2,* which is thought to promote GBM stem-like cells by activating EMP [72]. Finally, the SNAI1-IS module distinctively combined a triad of *LAG3, FOXP3,* and *SNAI1*, which pointed to dynamics of cell plasticity and immune evasion specific to oligodendroglioma. Modules significantly associated with astrocytomas (iTR-MES, AC-RA) had a strong negative correlation with oligodendrogliomas, confirming subtype-specific preferential residence in a more-mesenchymal state for the former and stem-like state for the latter.

Compared to the other two CG subtypes, undefined gliomas most strongly expressed the heterogeneous module 1 (G-MAL), which combined many genes associated with GBM malignancy and stemness. *ZEB1/2* were included in the G-MAL module, suggesting greater implication of these EMP transcription factors in undefined gliomas. However, regardless of neoplastic cell morphology, *ZEB1/2* were frequently upregulated across all CG subtypes by cells with cytoplasmic E-cadherin expression (Fig. 4d). Upregulation of the EPC-BR module, which combined *CD19, RORC, MMP2,* and *EPCAM*, also uniquely suggested more consistent interactions between B-cells and neoplastic cells upregulating *EPCAM* in undefined gliomas. Finally, module 4 (iMES-EX) and module 6 (TAN-TRF) were negatively correlated with oligodendrogliomas (*r* = -1.15), though were not strongly upregulated in either astrocytomas or undefined gliomas.

We then used Spearman’s rank correlation to quantify module co-occurrence across patients agnostic of CG subtype (Fig. 4b). The most significant correlations were found for G-MAL/iMES-EX (*r* = 0.75), G-MAL/SNAI1-IS (*r* = 0.74), G-MAL/OPC-like (*r* = 0.65), TAM-LEX/iMES-EX (*r* = 0.7), CTL-NK/CHK-T (*r* = 0.8), and TAN-TRF/CHK-T (*r* = 0.71). First, the co-occurrence of G-MAL with not only iMES-EX, but also SNAI1-IS and OPC-like supported the existence of heterogeneous niches across phenotypes, where generically malignant TME niches co-occur with specialized niches that leverage specific EMP and immune evasion mechanisms. As a direct reflection of this first observation, the strong correlation between TAM-LEX and iMES-EX was also in line with the standard interplay between EMP and immunosuppression, where more-mesenchymal neoplastic cells assemble a TAM-and exhausted T-cell-rich microenvironment. Although OPC-like niches heterogeneously co-occurred with G-MAL niches, there was a moderate to strong negative association between OPC-like niches and modules upregulated in astrocytomas (iTR-MES, MES-IL6, AC-RA).

There was a slight OPC-like/TAM-LEX negative correlate, echoing the invert correlate found in TAM-LEX and the CG oligodendroglioma subtype. We further investigated the position of each individual case for TAM-LEX/iMES-EX, TAM-LEX/iTR-MES, and TAM-LEX/TAN-TRF along our linear model to ensure the absence of outliers (Fig. 4e). Though astrocytoma and undefined glioma cases had a more heterogeneous distribution, each case scored high for at least one of the modules. Oligodendrogliomas, on the other hand, consistently had a low score on either axis, supporting a true tendency to lower occurrence of all tested modules.

Then, the co-occurrence of CTL-NK and CHK-T across all subtypes demonstrated that beyond sparse T-cell infiltration, CG also replicates the trend to immune exhaustion found in HG. Importantly, though CHK-T upregulation alone is not sufficient to support immune exhaustion, its co-occurrence with CTL-NK indicates that cytotoxic T-cells that penetrate the TME in CG become rapidly exhausted. This correlation provided a strong rationale for the lack of association found between CD3^+^ T-cell infiltration and OS in our cohort (Fig. 3b).

We completed our analysis by testing preferential module upregulation per CG subtype in intratumoral cores compared to the invasive front and “healthy” control tissue cores (Fig. 4f). Though no overt trend was found in undefined gliomas, astrocytomas and oligodendrogliomas each showed differential module upregulation. For astrocytomas, the AC-RA module was most strongly expressed at the invasive front, while the iMES-EX module was upregulated by intra-tumoral cores. By overlying the AC-RA module with tissue cores, we found that this most likely reflected AC-RA upregulation in cells resembling reactive astrocytes near the invasive front, suggesting an overlap between the phenotype of neoplastic ALC and reactive astrocytes. Preferential intra-tumoral upregulation of iMES-EX was compatible with the module’s prominent impaired Th17 response/immune dysfunction/exhaustion-related components (*BAG3, JAK1, IL17RA, HAVCR2, NFE2L2, STAT3, TCF7*). For oligodendrogliomas, TAM-LEX and CTL-NK were found more often in “healthy” control tissue, though data points were sparse and widely spread out for control cores.

All in all, these supported that regardless of subtype, CG demonstrates concurrent T-cell exhaustion and immunosuppressive macrophage polarization compounded by EMP activation in cancer cells. Specialized stem-like (OPC-like) and more-mesenchymal (iMES-EX, iTR-MES) niches with intrinsic features of immune evasion co-occurred with the G-MAL module across subtypes, reflecting intra-tumoral and inter-patient heterogeneity.

## Discussion

Our combined findings show that CG spatially recapitulates the heterogeneous EMP-immunosuppression niches previously described both by sequencing and tissue analysis in high-grade HG [18,61,73]. Tregs and exhausted T-cells co-localized with TAMs not only in PVC, but also in EMP-high niches. While each CG subtype relied on distinct EMP/immune evasion modules, associations between EMP/stemness and immune evasion were consistently found not only across all three CG subtypes, but also between patients.

Immunotherapy is increasingly considered a potential treatment option for both HG and CG. Anti-tumor vaccines, immune checkpoint inhibitors (anti-PD-1/PD-L1, anti-CD200), and CAR-T cells have been investigated in human and canine clinical trials, though these have failed to produce curative outcomes so far in either species [3,5,74–76]. Besides the uniquely inaccessible nature of the cerebral microenvironment, this lack of success can be explained by limited immune-mediated targeting of cancer cells. Failure has notably been attributed to a combination of sparse T-cell infiltration, widespread TME immunosuppression (Tregs, TAMs, exhausted T-cells), low target expression (PD-L1), and weak interferon signaling [27,74]. In human carcinomas, such as colorectal cancer, breast cancer, and squamous cell carcinoma, TME immunosuppression is known to correlate with EMP activation [16,77–80]. The heterogeneity of intratumoral EMP niches is thought to correlate with similarly heterogeneous immunotherapy response rates in patients [16,81].

Comparable dynamics have been proposed for non-canonically epithelial cancers, including melanoma and HG, with an emphasis on human GBM [4,11,12,18,20,23,81,82]. Recently, spatial transcriptomics analysis of human high-grade HG samples has likewise uncovered a trend toward assembling heterogeneous plastic niches, with markedly immunosuppressive microenvironments tracking along areas of necrosis and preferential perivascular distribution of OPC-like cells [73,83]. Low-grade HG are thought to exhibit similar cell-plasticity immunosuppression trends, though the mechanistic and spatial evidence is currently stronger in GBM [24,25].

A growing body of comparative veterinary research has explored cell plasticity and cell states as markers of immunosuppression and indicators of targetable therapeutic pathways [35,38,39,84]. However, although robust similarities between the biology of HG and CG have been established on multiple levels, no prior study has formally documented whether tumor heterogeneity correlates with immunosuppression in CG. Confirming that CG replicates these dynamics would strengthen the translational value of canine immunotherapy trials, with the advantage of potentially being run before or in parallel with human clinical trials. This is further reinforced by evidence of decreased OS combined with EMP downregulation following immunotherapy in French Bulldogs with high-grade CG, mirroring the shift away from a predominant MES-like phenotype in HG patients with acquired immunotherapy resistance [17,35]. Therefore, we aimed to determine whether CG spatially replicates the established correlations between cell plasticity and immunosuppression found in HG, with an emphasis on intratumoral heterogeneity.

We first focused on cancer cell morphology in relation to cell plasticity and stemness. Collectively, our results demonstrate the existence of heterogeneous neoplastic niches not only across patients but also within individual tumors. Though spindle-shaped UNC had a more-mesenchymal phenotype across CG subtypes, their protein and gene expression profiles varied. UNC and SLC were additionally associated with diverse EMP and stemness signatures across all three CG subtypes, supporting the idea that morphology alone is not sufficient to infer cell state. This is in line with HG literature and diagnostic guidelines where molecular identity and gene networks can vary across neoplastic cells and tumor types with shared histomorphological features [18,21,22,85]. UNC and SLC frequently displayed dual cytoplasmic E-cadherin and vimentin expression, a phenomenon usually described in hybrid E/M cells. While hybrid E/M states are well-documented in humans, evidence for comparable EMP features in cancers affecting domestic dogs and cats is only beginning to accumulate [36–38,86]. Among these features, dual expression of vimentin and cytoplasmic E-cadherin has been previously demonstrated in canine and feline mammary carcinomas exhibiting highly atypical cell morphology, upregulated EMP genes, and strong immunosuppressive characteristics [38,39]. A key cross-verification supporting true cytoplasmic E-cadherin expression rather than non-specific cross-labeling in our cohort is the consistent upregulation of *ZEB1/2* in tumor cores showing strong cytoplasmic E-cadherin labeling.

Under physiological conditions, *ZEB1* downregulates E-cadherin by directly repressing *CDH1* transcription. Consequently*, ZEB1* is often upregulated in cancers with EMP activation as cells become more motile and migratory. *ZEB1* upregulation has also been specifically linked to loss of membranous E-cadherin and abnormal cytoplasmic sequestration [69]. Concurrent upregulation of other EMP transcription factors (*SNAI1/2, TWIST1/2*) in the same TMA tissue cores further supports our interpretation that cytoplasmic E-cadherin-expressing cells have activated EMP pathways. Though E-cadherin is not expressed in the healthy brain, its upregulation has been reported in high-grade HG distinctly from other cadherins [87]. *CDH1*/E-cadherin is additionally known to be upregulated by OPCs and differentiating oligodendrocytes, which supports that cells with cytoplasmic E-cadherin-expressing cells residing in a de-differentiated, stem-like state [88]. High-grade HG that upregulate E-cadherin generally carry a worse prognosis, further reinforcing the parallel between EMP activation, invasiveness, and aggressive biological behavior [87].

With regard to markers of stemness evaluable by both IHC and transcriptomic analysis, tumor cores and cells that upregulated *OLIG2*/OLIG2 and EMP transcripts did not always upregulate *SOX10*/SOX10. Biologically, *OLIG2* is expressed earlier than *SOX10* in developing oligodendrocyte precursors and has been shown to directly regulate *SOX10* expression in mice [89]. Though mechanistic investigation would be required for confirmation, this raises the possibility that, regardless of subtype, CG with high *OLIG2*/OLIG2expression and low *SOX10*/SOX10 expression are more stem-like, thus potentially more immunosuppressive than other gliomas.

Subtype-specific EMP trends were also apparent. Astrocytomas displayed a tendency to acquiring a more-mesenchymal and invasive phenotype, notably supported by significant upregulation of the genes *TWIST1, PTX3, FOSL1,* and *GPRC5A*. Oligodendrogliomas instead resided in a more stem-like state, which was reflected in combined upregulation of *SOX4, SOX10*, and, together with undefined gliomas, of *PDGFRA*. A unique feature of oligodendrogliomas was differential upregulation of *ALDH1L2*, which sustains NADPH synthesis and redox homeostasis in the brain. Upregulation of *ALDH1L2* in HG is thought to stimulate sustained residence in a stem-like state by curbing oxidative damage, hence acting as a tumor-promoting factor [90].

Then, investigation of the immune TME revealed that intra-tumoral T-cells in all CG subtypes either had a regulatory phenotype or were exhausted, as evidenced by upregulation of lymphocytic immune checkpoints (*CTLA4*, *PDCD1*, *LAG3*, *HAVCR2*, *TIGIT*, *CD244*). Each CG subtype also hosted TAMs, with a similar baseline gene signature, further substantiating the immunosuppressive nature of EMP-enriched and/or stem-like neoplastic microenvironments in CG. T-cells and TAMs co-occurred spatially as clusters, which often aggregated in PVC and mapped preferentially to EMP-high niches. Although we did not identify subtype-specific upregulation of T-cell exhaustion markers, astrocytomas differentially upregulated *TGFB1*. TGF-β1 is a pleiotropic immune cytokine that is widely immunosuppressive and a major inducer of T-cell polarization toward a regulatory phenotype. Aside from its immunosuppressive functions, TGF-β1 is a pivotal inducer of EMP in cancer cells [91]. Thus, *TGFB1* upregulation in astrocytomas was consistent with their generally more-mesenchymal phenotype [91,92]. *GZMB* was also upregulated specifically in oligodendrogliomas compared, suggesting relatively increased recruitment of cytotoxic effector T-cells. However, based on our comparative *Smoothie* modules analysis of TMA cores sampled from the tumor center, invasive front, and surrounding neuroparenchyma, these are likely confined to the periphery of oligodendrogliomas and surrounding tissue, further limiting their anti-tumoral potential.

Finally, module-level *Smoothie* analysis highlighted two notable features. First, each subtype was associated with distinct gene modules linking cell plasticity and immunosuppression, which indicated subtype-intrinsic reliance on specific EMP and immune evasion programs.

Astrocytomas preferentially upregulated the TAM-LEX, iTR-MES, MES-IL6, and AC-RA modules, again highlighting strong parallels with EMP and immunosuppression dynamics found in other cancers. The MES-IL6 module was of particular interest; by combining *EGFR, IL6, VIM*, and *TWIST1*, this module supported more-mesenchymal differentiation potentially driven by IL-6 [53]. IL-6 is heavily secreted by reactive astrocytes in neuroinflammatory contexts, and is released in high amounts in high-grade HG, where it is thought to contribute to tumor progression and invasiveness via pro-tumorigenic IL-6/STAT3 signaling [93–97]. Aside from HG, IL-6 and TWIST1 engage in a feedback loop that mediates pancreatic cancer progression [71]. Together, these observations support an interplay between IL-6 and more-mesenchymal differentiation in canine astrocytomas. This may reflect sustained IL-6 secretion by both neoplastic and reactive astrocytes.

Besides residence in a pronounced OPC-like state, oligodendrogliomas uniquely upregulated a small network of 3 genes: *FOXP3*, *LAG3*, and *SNAI1*. *SNAI1* is a driver of more-mesenchymal cell states and is upregulated via TGF-β-mediated Smad3 signaling [98]. In addition, TGF-β is a canonical potent inducer of the transcription factor FoxP3 in CD4^+^ T-cells, thus leading to polarization toward a regulatory (immunosuppressive) phenotype [99]. Lag-3, which is an immune checkpoint protein, is found not only on effector T-cells, but also on FoxP3^+^Tregs. While considered a marker of exhaustion and dysfunction in effector T-cells, Lag-3 expression in FoxP3^+^Tregs is an active part of the immune response-dampening machinery of a subset Tregs [100]. Thus, preferential upregulation of this module by oligodendrogliomas may support subtype-specific enrichment in a subset of FoxP3^+^Lag-3^+^Tregs induced by microenvironments with high *SNAI1* upregulation. These findings suggest that astrocytomas may be better poised for investigating STAT3 inhibition, while oligodendrogliomas may be more useful in studying LAG-3 inhibition and/or targeting FoxP3^+^ Tregs as a therapeutic target.

Second, we identified correlations between these modules that held agnostic to CG subtype. This shows that even if EMP activation and immune cell exhaustion are important elements to take into consideration for planning care and assessing immunotherapy response, patient-intrinsic factors introduce a degree of variability that may not always be predictable from subtype alone. Accordingly, this suggests that CG is ideally placed to replicate the tendency to heterogeneous immunotherapy response seen in humans. Caution should be exercised, however, against directly conflating CG histological subtypes with HG tumor types, since CG subtyping does not carry the same molecular or therapeutic implications as in humans [30].

A key limitation of this study was the use of FFPE tissues, which precludes interventional experiments and may introduce minor artifacts resulting from formalin-induced epitope cross-linking. We mitigated this limitation by implementing *Smoothie*-based analysis to extract significantly relevant concurrent gene upregulation instead of relying solely on descriptive tissue analysis. We further argue that our approach demonstrates the value of archived canine FFPE tissues as an untapped source of biological data. Despite their generally being archived together with clinical metadata, such resources regrettably remain underused because of the lack of optimization of spatial -omics platforms for FFPE samples derived from non-human, non-rodent species. Additional limitations include the relatively small size of our cohort, though the use of TMA conversely increased our sample size significantly compared to traditional single tissue-based spatial transcriptomics analysis. Leveraging the use of TMA specifically allowed us to uncover inter-patient variability that would otherwise have been missed entirely. Though CG oligodendrogliomas included in this study were mostly high-grade tumors (9/10 cases), this bias directly results from the natural prevalence of high-grade oligodendrogliomas in the dog and is not expected to skew results [30,35]. An immediate next step would be to expand our limited 100-gene panel to better correlate cell phenotypes in CG and immunosuppression with the specific subtypes established by Neftel *et al*, which we were unable to compare our samples to given the incompatibility of FFPE with untargeted spatial transcriptomics approaches.

## Conclusions

We conclude that CG harbors heterogeneous stem-like/EMP immunosuppressive niches analogous to those described not only in high-grade HG, but also in human, feline, and canine carcinomas, where EMP activation correlates with T-cell exhaustion, recruitment of TAMs, and increased Treg polarization. Although the concept of EMP cannot be directly translated to HG/CG given their non-epithelial nature, neoplastic cells in HG/CG seem to rely on similar pathways to dynamically shift between cell states. These findings strongly position CG as a spontaneous, translationally relevant model that naturally integrates EMP-immunosuppression dynamics commonly associated with immunotherapy resistance. As immunotherapy becomes more widely available for canine patients, our results further indicate that each CG subtype may represent a distinct therapeutic challenge with varied potential for immunotherapy resistance, underscoring the importance to study CG subtype separately in translational research. Finally, our findings also support the value of improving ante-mortem subtyping and phenotyping methods for CG, which could help better identify strong immunotherapy candidates and strengthen the translational relevance of canine clinical trials.

## List of abbreviations

ALC: Astrocyte-like cells
CG: Canine glioma
CNPase: 2’,3’-Cyclic-nucleotide 3’-phosphodiesterase
CNS: Central nervous system
E-cad: E-cadherin
EMP: Epithelial-mesenchymal plasticity
FFPE: Formalin-fixed, paraffin-embedded
GBM: Glioblastoma
GFAP: Glial fibrillary acidic protein
HG: Human glioma
ICI: Immune checkpoint inhibitors
MP: Glomeruloid microvascular proliferation
OLC: Oligodendrocyte progenitor-like cells
OLIG2: Oligodendrocyte transcription factor 2
OS: Overall survival
Rac: Reactive astrocytes
SLC: Stem-like cells
SOX10: SRY (sex determining region Y)-box 2
SOX2: SRY (sex determining region Y)-box 2
TAM: Tumor-associated macrophage
TMA: Tissue micro-array
TME: Tumor microenvironment
Treg: Regulatory T-cell
UNC: Undifferentiated cells
Vim: Vimentin

## Declarations

### Ethics approval and consent to participate

All samples included in this study were FFPE canine samples obtained from the archives of the New York State Animal Health Diagnostic Center and the Cornell University Neuropathology Service. All samples included in this study were submitted to the AHDC from a referring veterinarian for diagnostic purposes. Per internal protocol, all samples received by the NSVDL are the property of the NYSVDL and do not need owner consent.

### Consent for publication

Not applicable.

### Availability of data and material

All data supporting the findings of this study are available within the paper and its Supplementary Information.

### Competing interests

The authors declare that they have no competing interests.

### Funding

No external funding was received for this study.

### Competing interests

The authors declare that they have no competing interests.

### Authors’ contributions

EAD and SRN established the experimental design and selected relevant samples. SRN conducted the digital image analysis and descriptive omics analysis and assembled related panels. ADM and EAD diagnosed and graded canine glioma samples. ENJ, JH, and GS diagnosed canine patients in the clinic and provided patient samples and relevant history. AS, IV, and PS performed the computational transcriptomics data analysis, and AS assembled related panels. All authors read and approved the final manuscript.

## Acknowledgments

The authors acknowledge all staff of the histology laboratory of the Cornell University Animal Health Diagnostic Center, for their invaluable support and assistance with tissue processing and staining; the 10x Genomics members Chengwei Zhong and Ivy Zhong, for providing technical support and helpful discussion; and the University of Rochester Medical Center’s Genomics Research Center, especially Jeffrey Malik, PhD, Elizabeth Pritchett, PhD, and Phillip Spinelli for their assistance with running the Xenium v1 analysis.

---

**Supplemental Figure 1.**
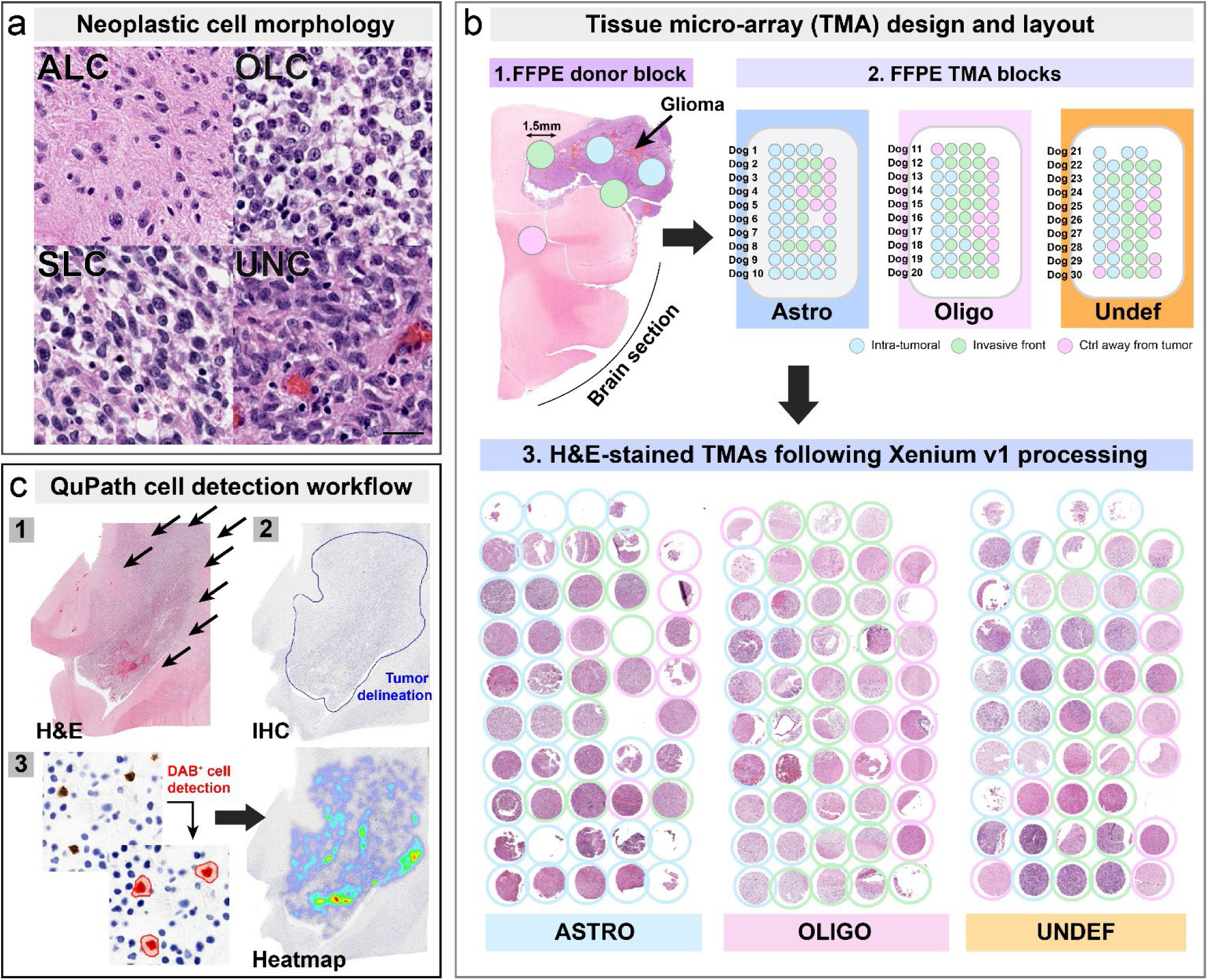
Experimental design. **(a) Neoplastic cell morphology**. Cells were categorized into astrocyte-like (ALC), oligodendrocyte-like (OLC), stem-like (SLC), and undefined (UNC). Scale bar: 25μm. **(b) Tissue micro-arrays (TMAs) design and preparation.** One block was designed per canine glioma subtype, each holding 5 tissue cores from 10 cases. The first row had 4 tissue cores only for orientation purpose; case 6 and case 28 each had 4 cores only as 1 tissue core was too small to be properly integrated for each case. TMAs were stained with hematoxylin and eosin (H&E) immediately after Xenium v1 analysis for direct overlap of gene clusters with histomorphological features. **(c)** QuPath cell detection workflow for CD3^+^ T-cell quantification and heatmap generation.

**Supplemental Figure 2.**
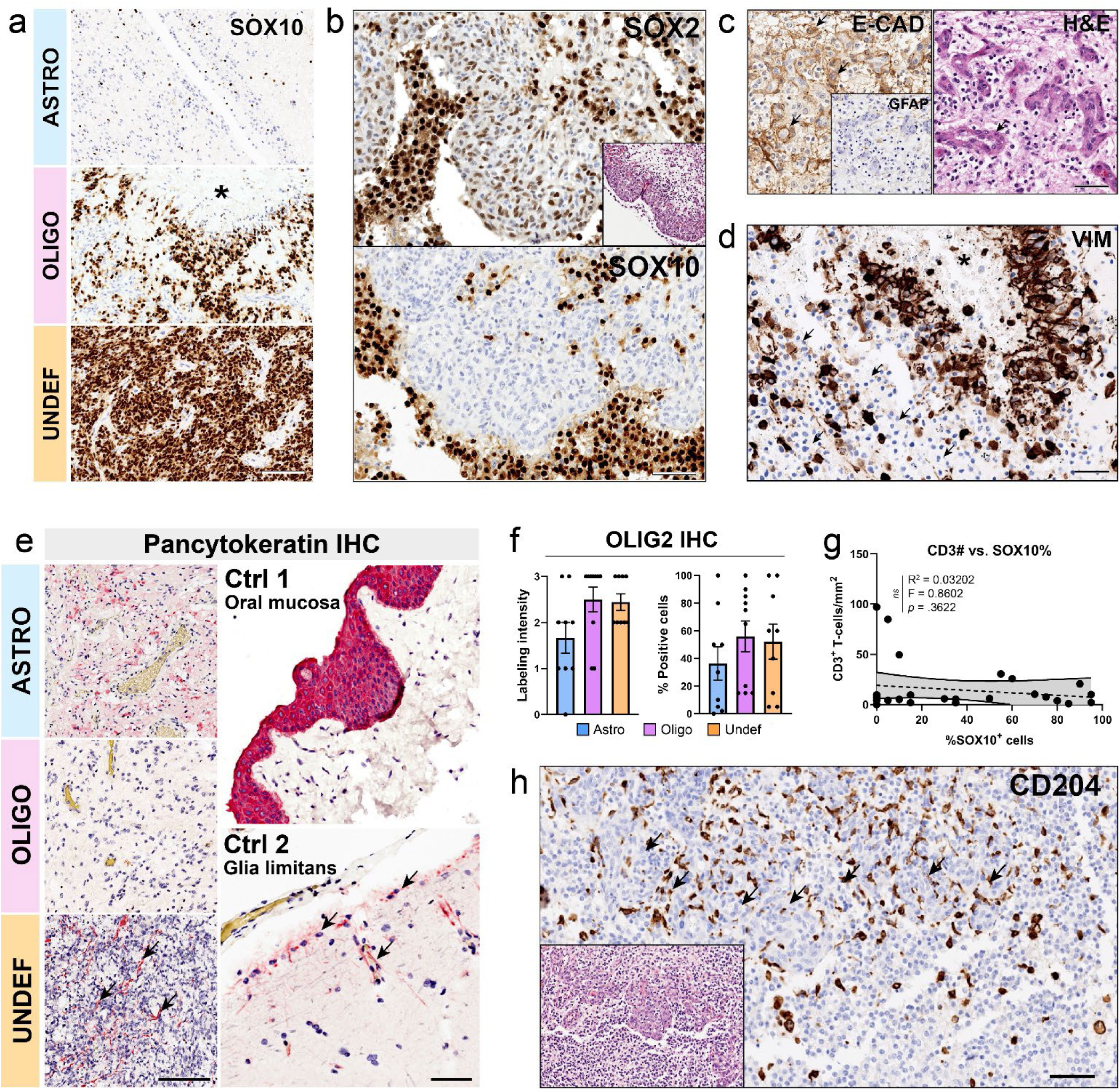
Cell plasticity and stemness IHC, and correlation with immune microenvironment. **(a)** In tumors with low or heterogeneous expression, SOX10 labeling was often found in sparse scattered and preferentially near the invasive front (ASTRO). SOX10 labeling was strong in pseudopalisading neoplastic cells (OLIGO). Undefined gliomas most often had extensive labeling of strong intensity (UNDEF). *: necrosis. Scale bar: 100 μm. **(b)** MPs had faint to moderate SOX2 labeling but lacked SOX10 labeling. Inset: H&E. Scale bar: 50 μm. **(c)** Atypical UNC cells surrounding proliferative vascular structures often had mixed membrane and cytoplasmic E-cadherin labeling. Inset: No cross-labeling with GFAP was observed. Scale bar: 25 μm. **(d)** Pseudopalisading neoplastic cells had strong vimentin labeling. Well-differentiated OLC lacked vimentin labeling (arrows). *: necrosis. Scale bar: 50 μm. **(e)** Though pancytokeratin labeling occurred in some tumors, the labeling pattern was redundant with GFAP (UNDEF, arrows). Scale bar: 100μm. The glia limitans exhibited aberrant pancytokeratin labeling in a control healthy canine brain, suggesting non-specific cross-reactive labeling (arrows). Scale bar: 25 μm. **(f)** OLIG2 IHC labeling was most extensive and intense in oligodendrogliomas and undefined gliomas. **(g)** No direct correlation between %SOX10-positive cells and CD3 T-cell infiltration was seen. **(h)** CD204+ cells preferentially gathered along MPs (arrows) and areas of necrosis. Scale bar: 50 μm.

**Supplemental Figure 3.**
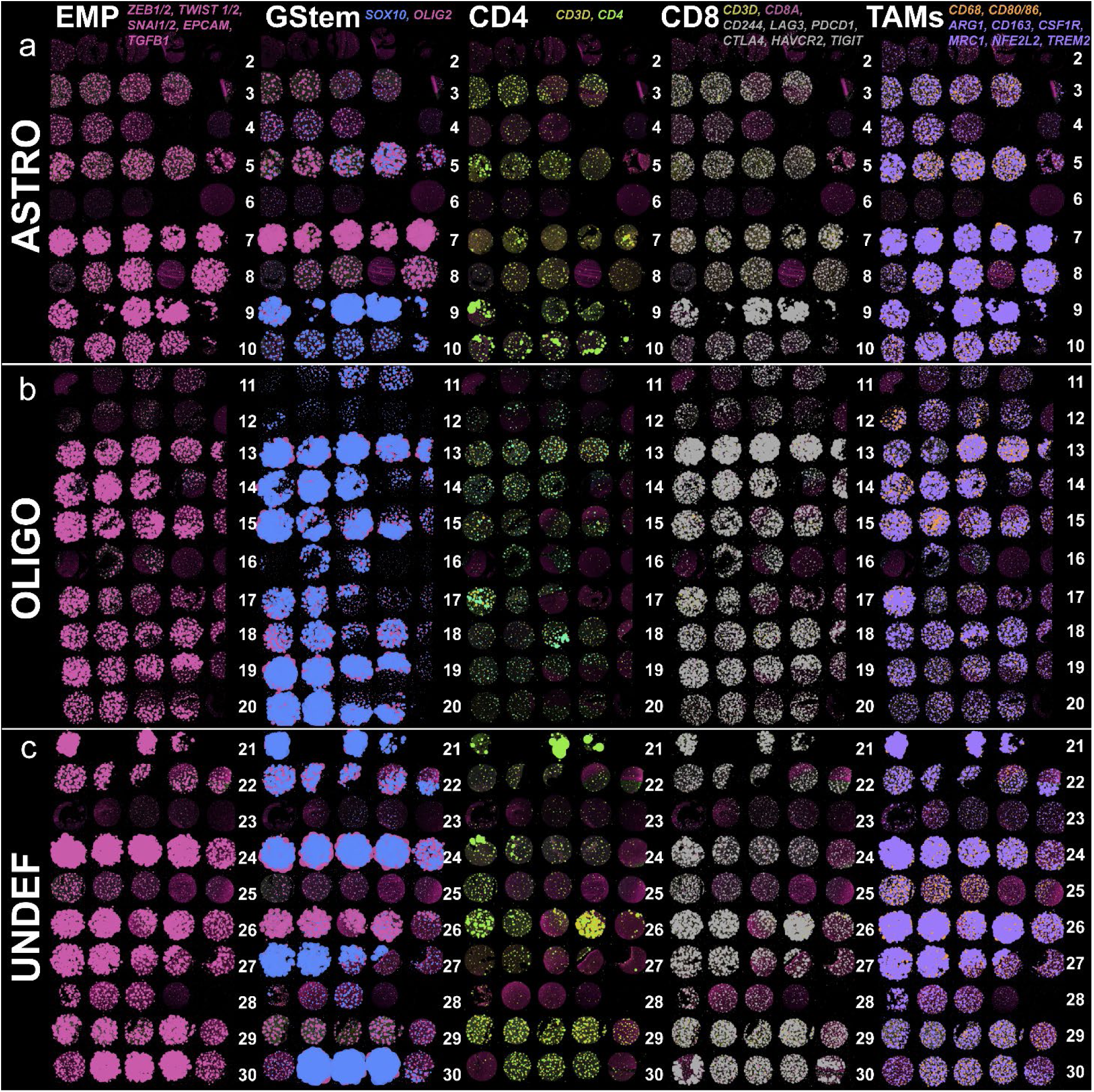
Spatial correlation between cell plasticity (EMP), glioma stemness (*SOX10*, *OLIG2*), CD4 T-cells (CD4), exhausted CD8 T-cells (CD8), and immunosuppressive macrophages (TAMs). **(a-c)** In all glioma subtypes, EMP transcripts upregulation correlated with CD8 T-cell exhaustion and TAMs infiltration. CD4 T-cells were also more numerous in these cores. Upregulated *SOX10* transcripts often correlated with upregulated EMP transcripts; oligodendroglioma was the only subtype for which *SOX10* and EMP transcript upregulation was consistently proportional. White numbers: case numbers (1 case/row). Astrocytoma: rows 2-10, Oligodendroglioma: rows 11-20, Undefined glioma: rows 21-30. Case 1 was discarded at quality check.

## Notes

### Competing Interest Statement

The authors have declared no competing interest.

